# Thermophilic carboxylesterases from hydrothermal vents of the volcanic island of Ischia active on synthetic and biobased polymers and mycotoxins

**DOI:** 10.1101/2022.09.17.508236

**Authors:** Marco A. Distaso, Tatyana N. Chernikova, Rafael Bargiela, Cristina Coscolín, Peter Stogios, Jose L. Gonzalez-Alfonso, Sofia Lemak, Anna N. Khusnutdinova, Francisco J. Plou, Elena Evdokimova, Alexei Savchenko, Evgenii A. Lunev, Michail M. Yakimov, Olga V. Golyshina, Manuel Ferrer, Alexander F. Yakunin, Peter N. Golyshin

**Author notes:** Corresponding authors at: the Centre for Environmental Biotechnology, Bangor University, LL57 2UW Bangor, UK (P.N. Golyshin); ICP, CSIC, Marie Curie 2, 28049 Madrid, Spain (M. Ferrer). E-mail addresses (M. Ferrer), (P.N. Golyshin) (M. Ferrer), (P.N. Golyshin). Equal contributions (Manuel Ferrer, Alexander F. Yakunin, Peter N. Golyshin).

## Abstract

Hydrothermal vents have a widespread geographical distribution and are of high interest for investigating microbial communities and robust enzymes for various industrial applications. We examined microbial communities and carboxylesterases of two terrestrial hydrothermal vents of the volcanic island of Ischia (Italy) predominantly composed of Firmicutes (*Geobacillus* and *Brevibacillus* spp.), *Proteobacteria* and *Bacteroidota*. High-temperature enrichment cultures with the polyester plastics polyhydroxybutyrate (PHB) and polylactic acid (PLA) resulted in an increase of *Thermus* and *Geobacillus* spp., and to some extent, *Fontimonas* and *Schleiferia* spp. The screening at 37-70ºC of metagenomic fosmid library from above enrichment cultures resulted in identification and successful production in *Escherichia coli* of three hydrolases (IS10, IS11 and IS12), all derived from yet uncultured Chloroflexota and showing low sequence identity (33-56%) to characterized enzymes. Enzymes exhibited maximal esterase activity at temperatures 70-90ºC, with IS11 showing the highest thermostability (90% activity after 20 min incubation at 80ºC). IS10 and IS12 were highly substrate-promiscuous and hydrolysed all 51 monoester substrates tested. Enzymes were active with polyesters (PLA and polyethylene terephthalate model substrate, 3PET) and mycotoxin T-2 (IS12). IS10 and IS12 had a classical α/β hydrolase core domain with a serine hydrolase catalytic triad (Ser155, His280, and Asp250) in the hydrophobic active sites. The crystal structure of IS11 resolved at 2.92 Å revealed the presence of the N-terminal β-lactamase-like domain and C-terminal lipocalin domain. The catalytic cleft of IS11 includes catalytic residues Ser68, Lys71, Tyr160, and Asn162, whereas the lipocalin domain encloses the catalytic cleft like a lid contributing to substrate binding. Thus, this study has identified novel thermotolerant carboxylesterases with a broad substrate range including polyesters and mycotoxins for potential applications in biotechnology.

**IMPORTANCE:** High-temperature-active microbial enzymes are important biocatalysts for many industrial applications including recycling of synthetic and biobased polyesters increasingly used in textiles, fibres, coatings and adhesives. Here, we have discovered three novel thermotolerant carboxylesterases (IS10, IS11 and IS12) from high-temperature enrichment cultures from the Ischia hydrothermal vents incubated with biobased polymers. The identified metagenomic enzymes originated from uncultured Chloroflexota and showed low sequence similarity to known carboxylesterases. Active sites of IS10 and IS12 had the largest “effective volumes” among the characterized prokaryotic carboxylesterases and exhibited high substrate promiscuity, including hydrolysis of polyesters and mycotoxin T-2 (IS12). Though less promiscuous compared to IS10 and IS12, IS11 had a higher thermostability with high temperature optimum (80-90 ºC) for activity, hydrolysed polyesters, and its crystal structure revealed an unusual lipocalin domain likely involved in substrate binding. The polyesterase activity in these enzymes makes them attractive candidates for further optimisation and potential application in plastics recycling.

## INTRODUCTION

Environmental microbial communities and microorganisms represent an enormous reserve of biochemical diversity and enzymes for fundamental research and applications in biotechnology (1,2). However, the vast majority of environmental microbes have never been grown and characterised in the laboratory (3,4). The metagenomic approach has emerged as a strategic way to study unculturable microorganisms and their enzymes using various computational and experimental methods (5-7). Metagenomics includes shotgun sequencing of microbial DNA purified from a selected environment, high-throughput screening of metagenomic expression libraries (functional metagenomics), profiling of RNAs and proteins produced by a microbial community (meta-transcriptomics and meta-proteomics), and identification of metabolites and metabolic networks of a microbial community (meta-metabolomics) (8). Global DNA sequencing efforts and several large-scale metagenome sampling projects revealed the vast sequence diversity in environmental metagenomes and microbial genomes, as well as the presence of numerous unknown or poorly characterised genes (9-12). For example, a high-throughput project focused on carbohydrate-active enzymes has identified over 27,000 related genes and demonstrated the presence of glycoside hydrolase activity in 51 out of 90 tested proteins (13). Other large scale metagenomic projects include the Sargasso Sea sampling (over one million new genes discovered), the Global Ocean Survey (over six million genes), and human gut microbiome (over three million genes) (9-12). Thus, through the advent of metagenomics, we are starting to generate insights into the rich microbial worlds thriving in different environments. Nevertheless, a recent analysis of metagenome screening studies suggested that all representative types of environmental habitats (terrestrial, marine, and freshwater) are under-sampled and under-investigated (14). It is estimated that total number of microbial cells is 10^30^, whereas the natural protein universe exceeds 10^12^ proteins indicating that our knowledge of proteins and biochemical diversity on Earth is very limited (15-17). Therefore, the determination of protein function or enzyme activity for millions of genes of unknown function and biochemically uncharacterised proteins represents one of the main challenges of the postgenomic biology.

The approaches of experimental metagenomics include meta-transcriptomics, meta-proteomics, metabolomics, and enzyme screening (6,7,17-19). Activity-based screening of metagenome gene libraries represents a direct way for tapping into the metagenomic resource of novel enzymes. This approach involves expressing genes from metagenomic DNA fragments in *Escherichia coli* cells and assaying libraries of clones on agar plates for enzymatic activities using chromogenic or insoluble substrates (18). Importantly, this approach offers the possibility to identify novel families of enzymes with no sequence similarity to known enzymes. Screening of metagenome gene libraries from different terrestrial, marine, and freshwater environments has already expanded the number of new enzymes including novel nitrilases, glycoside hydrolases, carboxyl esterases, and laccases (14,20,21).

Carboxylesterases (EC 3.1.1.1) are a diverse group of hydrolytic enzymes catalyzing the cleavage and formation of ester bonds, which represent the third largest group of industrial biocatalysts (after amylases and proteases). Many esterases show a wide substrate range and high regio- and stereo-selectivity making them attractive biocatalysts for applications in pharmaceutical, cosmetic, detergent, food, textile, paper and biodiesel industries (22,23). Most of known carboxylesterases belong to the large protein superfamilies of α/β hydrolases and β-lactamases and have been classified into 16 families based on sequence analysis (22,24,25). A significant number of these enzymes have been characterised both biochemically and structurally, because they are of high interest for biotechnological applications (22,23,26). Screening of metagenome gene libraries and genome mining has greatly expanded the number of novel carboxylesterases including enzymes active against aryl esters or polymeric esters (polyesterases) (21-23,26,27). However, the increasing demand for environmentally friendly industrial processes has stimulated research on the discovery of new enzymes and their application as biocatalysts to meet the challenges of a circular bioeconomy (28,29). The global enzyme market is expected to grow from $8.18 billion in 2015 to $17.50 billion by 2024 (28). However, the majority of known enzymes are originated from mesophilic organisms, which have limited stability under harsh industrial conditions including high temperatures, extreme pH, solvents, and salts (30,31). Thus, the discovery of robust enzymes including carboxylesterases and engineering of more active variants represent the key challenges for the development of future biocatalytic processes. Extremophilic microorganisms are an attractive source of industrial biocatalysts, because they evolved robust enzymes that function under extreme conditions (high/low temperatures, high/low pH, salts) (14,26,30,32). In addition, extremophilic enzymes found in one environment are typically also tolerant to other extreme conditions making them attractive biocatalysts for various applications including depolymerization of natural and synthetic polymers (32-35).

Hydrothermal vents are extreme environments located in tectonically active sites, which are classified as terrestrial and marine (deep-sea and shallow-sea) systems (36). Hydrothermal vents are characterised by harsh physico-chemical conditions (high temperature and low pH) and are known as source of thermophilic microbes and enzymes with biotechnological importance. Although terrestrial hydrothermal vents have relatively easy access, they remain under-investigated compared to (sub)marine hydrothermal vents. To provide insights into microbial diversity of terrestrial hydrothermal vents, we analysed the natural microbial communities of two thermophilic hydrothermal vents located on the volcanic island of Ischia (Italy), as well as the effect of polyester plastic addition on these microbial communities using barcoded DNA sequencing of extracted DNA. Using activity-based metagenomic approach, we screened fosmid libraries for carboxylesterase activity using tributyrine agar plates, identified 14 unique fosmids encoding putative hydrolases, from which three soluble carboxylesterases (IS10, IS11, and IS12) were recombinantly produced in *E. coli* and biochemically characterised including substrate range and stability using both monoester and polyester substrates. The crystal structure of IS11 was resolved to reveal the N-terminal β-lactamase-like serine hydrolase domain connected to the C-terminal lipocalin domain. The active site of IS11 accommodated the conserved catalytic residues Ser68, Lys71, Tyr160, and Asn162, as well as numerous hydrophobic residues potentially involved in substrate binding. Structural models of IS10 and IS12 revealed classical α/β hydrolase domains with a catalytic serine hydrolase triad (Ser155, His280, Asp250), multiple hydrophobic residues in their active sites with the largest “effective volumes” reported for prokaryotic carboxylesterases.

## MATERIALS AND METHODS

### Environmental sampling sites and enrichment cultures

Sediment samples with water were collected in September 2018 from the geothermal areas of the volcanic island of Ischia (the Gulf of Naples, Italy). The samples were taken from the Cavascura hydrothermal springs (40.70403 13.90502): IS1 (pH 8.5, 45ºC) and IS2 (pH 7.0, 55ºC) and from the sandy fumaroles of Maronti beach near St Angelo (40.70101 13.89837): IS3 (pH 4.5, 75ºC) and IS4 (pH 5.0, 75ºC). For each sample, triplicate enrichment cultures were established containing different polymers or plastics as substrates, polylactic acid film (PLA, poly-D,L-lactide, M_w_ 10,000-18,000 Da), PLA polyhydroxybutyrate (PHB) and a commercial compostable polyester blend (P3, Blend) were kindly provided by the Biocomposites Centre, Bangor University, UK. Plastic films were cut (3 mm x 20 mm), washed in 70% ethanol and air-dried before adding to samples. For IS1 and IS2 cultures, modified DSMZ medium 1374 (https://bacmedia.dsmz.de/medium/1374) was used, which contained (g L^-1^): NaCl, 1; MgCl_2_.6H_2_O, 0.4; KCl, 0.1; NH_4_Cl, 0.25; KH_2_PO_4_, 0.2; Na_2_SO_4_, 4; NaHCO_3_, 0.1; CaCl_2_ .2H_2_O, 0.5. The medium was adjusted to pH 7.5 with 10N NaOH. For IS3 and IS4 cultures, modified DSMZ medium 88 (https://bacmedia.dsmz.de/medium/88) was used, which contained (g L^-1^): (NH_4_)_2_SO_4_, 1.3; KH_2_PO_4_, 0.28; MgSO_4_.7H_2_O, 0.25; CaCl_2_.2H_2_O, 0.07. The medium was adjusted to pH 4.5 with 10N H_2_SO_4_. Additionally, the trace element solution SL-10 (from DSMZ medium 320 https://bacmedia.dsmz.de/medium/320) was added at 1:1000 (vol/vol) to both media. Enrichment cultures contained 0.5 g of sample sediment and 0.25 g of a polymer in 10 mL of growth medium. The cultures were incubated at 50 °C (IS1-IS2) or 75 °C (IS3-IS4) with slow agitation (30 rpm) for 4 days, then culture aliquots (20% of the volume of enrichment cultures) were transferred to a fresh medium and incubated for 11 days under the same conditions (Table S1).

### DNA extraction and 16S rRNA amplicon sequencing

Prior to DNA extraction, the enrichment cultures (9 mL each) were vortexed and biomass was collected by centrifugation at 10,000 rcf for 10 min at 4°C. The pellets were resuspended in 250 μL of sterile phosphate-buffered saline (PBS, pH 7.5) and transferred to 1.5 mL tubes. High molecular weight DNA was obtained using the ZymoBIOMICS DNA Miniprep Kit (Zymo Research, Irvine, Ca, USA) in accordance with manufacturer’s instructions. Finally, DNA was eluted with 50 μL of nuclease free water. The quality of extracted DNA was assessed by gel electrophoresis, and DNA concentration was estimated using Qubit™ 4.0 Fluorometer dsDNA BR Assay Kit (Life Technologies, USA). The Illumina-compatible libraries of hypervariable V4 region of 16S rRNA gene were prepared by single PCR with dual-indexing primer system with heterogeneity spacer as described previously (37). Modified forward primer F515 (5’-GTGBCAGCMGCCGCGGTAA-3’) and reverse R806 prokaryotic primer (5’-GGACTACHVGGGTWTCTAAT-3’) were used. PCR reactions were performed using MyTaq™ Red DNA Polymerase (Bioline) in a Bio-Rad^®^ thermocycler with the following program: 95 °C for 2 min for denaturation followed by 30 cycles at 95 °C for 45 s, 50 °C for 60 s, 72 °C for 30 s, with a final elongation at 72 °C for 3 min. PCR products of approximately 440 bp were visualised by gel electrophoresis and gel-purified using the QIAEX II Gel Extraction Kit^®^ (QIAGEN). The purified barcoded amplicons were quantified by Qubit™ dsDNA BR Assay Kit (Life Technologies, USA), pooled in equimolar amounts and sequenced on Illumina MiSeq™ platform (Illumina Inc., San Diego, CA, USA) using paired-end 250 bp reads at the Centre for Environmental Biotechnology (Bangor, UK). Sequencing reads were processed and analysed as previously described (38). All statistical analysis was conducted using R programming environment (39) *prcomp* function and in-house scripts for graphical design.

### Preparation of the Ischia metagenome library from polyester enrichment cultures

High molecular weight DNA extracted from all enrichment cultures was combined in equimolar amounts and used to prepare two metagenomic fosmid libraries ‘IS_Lib1’ (Cavascura enrichments) and ‘IS_Lib2’ (Maronti enrichments) using the CopyControl™ Fosmid Library pCC2FOS Production Kit (Epicentre Technologies, Madison, USA). DNA was end-repaired to generate blunt-ended 5’-phosphorylated fragments according to manufacturer’s instructions. Subsequently, DNA fragments in the range of 30-40 kbp were resolved by gel electrophoresis (2 V cm^-1^ overnight at 4 °C) and recovered from 1% low melting point agarose gel using GELase 50X buffer and GELase enzyme (Epicentre). Nucleic acid fragments were then ligated to the linearized CopyControl pCC2FOS vector following the manufacturer’s instructions. After the in vitro packaging into the phage lambda (MaxPlax™ Lambda Packaging Extract, Epicentre), the transfected phage T1-resistant EPI300™-T1R *E. coli* cells were spread on Luria-Bertani (LB) agar medium containing 12.5 μg/ml chloramphenicol and incubated at 37 °C overnight to determine the titre of the phage particles. The resulting library had estimated titre of 14×10^4^ and 1×10^4^ non-redundant fosmid clones in IS_Lib1 and IS_Lib2 libraries, correspondingly. For long-term storage, *E. coli* colonies were washed off from the agar surface using liquid LB medium containing 20% (v/v) sterile glycerol and the aliquots were stored at −80 °C.

### Activity-based screening of the polyester enrichment metagenome library for esterase activity

The metagenomic library IS_Lib2 was screened for carboxylesterase/lipase activity as follows. The fosmid library was grown on LB agar plates containing 12.5 μg/ml chloramphenicol at 37°C overnight to yield single colonies. Then, 3,456 clones were arrayed in 9 × 384-well microtitre plates and cultivated at 37 °C in LB medium supplemented with 12.5 μg/ml chloramphenicol. After overnight growth, replica plating was used to transfer the clones onto the surface of large LB agar square plates (245 mm x 245 mm) containing 12.5 μg/ml chloramphenicol, 2 ml/L fosmid autoinduction solution (Epicentre), each plate supplemented with 0.3% (v/v) tributyrin (Sigma-Aldrich, Gillingham, UK). The original microtitre plates were stored at −80 °C with the addition of 20% (vol/vol) glycerol to enable the isolation of positive clones after the functional screenings. After an initial overnight growth at 37°C, the LB agar plates were incubated for 48 hours at 37, 50 or 70 °C. Positive hits were confirmed by re-testing of the corresponding fosmid clones taken from the original microtitre plate.

### Sequencing and analysis of metagenomic fragments

Positive fosmid clones were cultivated in 100 mL LB medium containing 12.5 μg/ml chloramphenicol and 2 ml/L fosmid autoinduction solution (Epicentre) at 37°C overnight. Biomass was collected by centrifuging at 5,000 rpm for 30 min and fosmid DNA was extracted from the pellet using the QIAGEN Plasmid Midi Kit (QIAGEN) following the manufacturer’s instructions. Approximate size of the cloned fragments was assessed on agarose gel electrophoresis after double endonuclease digestion with *Xba*I and *Xho*I (New England Biolabs, Ipswich, MA, USA). The Sanger sequencing of the termini of inserted metagenomic fragments of each purified fosmid was done at Macrogen Ltd. (Amsterdam, The Netherlands) using standard pCC2FOS sequencing primers (Epicentre). Non-redundant fosmids were selected, their DNA concentrations were quantified by Qubit™ 4.0 Fluorometer dsDNA BR Assay Kit (Invitrogen), pooled in equimolar amounts and prepared for Illumina MiSeq® sequencing. Pooled DNA was fragmented using the Bioruptor Pico Sonicator (Diagenode, Denville, NJ, USA) with parameters adjusted to obtain 400-600 bp fragments. The fragment library was prepared using the NebNext Ultra II DNA Library preparation kit (New England Biolabs, Ipswich, MA, USA) according to the manufacturer’s instructions. The obtained library was sequenced on MiSeq® platform (Illumina, San Diego, USA) using a microflow cell 300-cycles V2 sequencing kit. Obtained paired end reads were subjected to quality filtering, trimming and assembly as previously described (40). Gene prediction and primary functional annotation were performed using the MetaGeneMark annotation software (http://opal.biology.gatech.edu) (41). Translated protein sequences were annotated using BLAST searches of UniProt and the non-redundant GenBank databases (42). Multiple sequence alignments were generated using MUSCLE application (43) and visualised on Geneious v.9 (Biomatters, New Zealand). The Neighbour-Joining and maximum likelihood trees were constructed in MEGA X (44) using the settings for the Poisson model and homogenous patterning between lineages. The bootstrapping was performed with 1,000 pseudoreplicates.

### Gene cloning, expression and purification of selected proteins

Selected gene candidates were amplified by PCR in a T100 Thermal Cycler (Bio-Rad) using Herculase II Fusion Enzyme (Agilent, Cheadle, UK) with oligonucleotide primer pairs incorporating pET-46 Ek/LIC vector adapters (Merck, Darmstadt, Germany). PCR products were then purified and cloned into the above pET-46 Ek/LIC vectorharbouring an N-terminal 6xHis tag, as described by the manufacturer. The DNA inserts in the resulting plasmids were verified by Sanger sequencing at Macrogen Ltd. (Amsterdam, The Netherlands) and then transformed into *E. coli* BL21(DE3) for recombinant protein expression. *E. coli* BL21(DE3) cultures harbouring pET-46 Ek/LIC were grown on LB medium to mid-log growth phase (OD_600_ 0.7-0.8), induced with isopropyl-β-d-thiogalactopyranoside (IPTG, 0.5 mM) and incubated at 20°C overnight. Cells were disrupted by sonication as reported earlier (45) and recombinant proteins were purified using metal-chelate affinity chromatography on Ni-NTA His-bind columns. Protein size and purity were assessed using denaturing gel electrophoresis (SDS-PAGE), and protein concentration was measured by Bradford assay (Merck, Gillingham, UK).

### Enzyme assays

Carboxyl esterase activity of purified proteins against ***p*-nitrophenyl (*p*NP) or** α**-naphthyl (**α**N) esters** was determined by measuring the amount of *p*-nitrophenol released by esterase-catalysed hydrolysis essentially as described previously (27,45). Under standard assay conditions the reaction mixture contained 50 mM potassium phosphate buffer (pH 7.0), 1 mM *p*-nitrophenyl butyrate as substrate, and 0.2-1.8 μg of enzyme in a final volume of 200 μl. Reactions were incubated at 30°C for 3-5 min and monitored at 410 nm (for *p*NP esters) or 310 nm (for αN esters). Non enzymatic hydrolysis of ester substrates was subtracted using a blank reaction with denatured enzyme. The effect of pH on esterase activity was evaluated using the following buffers: sodium citrate (pH 4.0 and 5.0), potassium phosphate (pH 6.0 and 7.0), Tris-HCl (pH 8.0 and 9.0). The activity was monitored at 348 nm (the pH-independent isosbestic wavelength of *p*-nitrophenol). The effect of temperature on esterase activity was studied using a range of temperatures (from 20°C to 95°C). In order to assess the thermal stability of purified esterases, the enzymes were dissolved in potassium phosphate buffer (pH 7.0) and preincubated at the indicated temperature for 20 min. The enzyme solutions were then cooled down on ice and the residual activity was measured under standard conditions (at 30 ºC). Substrate specificity of purified enzymes was analysed using model *p*NP- and αN-esters with different chain lengths: *p*NP-acetate (C2), αN-propionate (C3), *p*NP-butyrate (C4), αN-butyrate (C4), *p*NP-hexanoate (C6), *p*NP-dodecanoate (C12), and *p*NP-palmitate (C16), obtained from Sigma-Aldrich and Tokyo Chemical Industry TCI. Kinetic parameters for these substrates were determined over a range of substrate concentrations (0.012-4 mM; 30 ºC) and calculated by non-linear regression analysis of raw data fit to Michaelis-Menten function using GraphPad Prism software v.6. Hydrolysis of 44 **soluble non-chromogenic monoester substrates** (Table S2) and **T-2 mycotoxin** (Merck Life Science S.L.U., Madrid, Spain) was assayed at 37ºC using a pH indicator assay with Phenol Red and monitored at 550 nm (46). The reaction products of enzymatic degradation of T-2 mycotoxin were analysed using reversed phase chromatography on a Waters 600 HPLC system equipped with a Zorbax Eclipse Plus C18 column (Agilent, 4.6 × 100 mm, 3.5 μm, 40ºC) and a light scattering detector (ELSD). The reaction products were separated using gradient elution (1.0 ml/min) with acetonitrile (with 0.2% (vol/vol) formic acid) and water (5%: 1 min, 5%-95%: 9 min, 95%: 3 min, 5%: 7 min). Polyester depolymerization activity of purified proteins against 3PET (bis(benzoyloxyethyl) terephthalate) was measured using 1.5% agarose plates containing 0.2% of emulsified polyesters. 3PET was purchased from CanSyn Chem. Corp. (Toronto, Canada). Agarose plates with emulsified 3PET were prepared as described previously (47). After protein loading, the plates were sealed and incubated at 37 ºC for 1-5 days. The presence of polyesterase activity was indicated by the formation of a clear zone around the wells with proteins. Apart from plate assays, activity assays of IS10, IS11 and IS12 for **3PET suspension hydrolysis** were performed in 50 mM Tris-HCl buffer, pH 8.0, at 30 °C, in a shaker 600 at rpm, the final reaction volume for each experiment was 0.2 mL, and the final protein amount 50 μg. The reactions were terminated after 13 h by filtering reaction mixture on a 10 kDa spin filter. 10 μL of filtrate was analysed using the high-performance liquid chromatography system (HPLC), Schimadzu, Prominence-I (Milton Keynes, UK) equipped with a Schimadzu C18 Shim-pack column (4.6 × 150 mm, 5 μm). The mobile phase was 25 % (vol/vol) methanol with 0.1% (vol/vol) H_3_PO_4_ in HPLC-grade water at a flow rate of 0.7 mL min^-1^ for 2 min, following increase to 55 % of methanol to 118 min, followed by 25% methanol at 22 min; the effluent was monitored at the wavelength of 240 nm, the column was conditioned at 40 ºC. The hydrolytic products of mono(2-hydroxyethyl)terephthalic acid (MHET), bis(2-hydroxyethyl)terephthalate BHET and terephthalic acid (TPA) were identified by comparing the retention times with their standards, reactions without enzyme were served as negative controls. All samples of each experiment were analysed in triplicate. **Enzymatic activity against PLA** was assayed by measurement of lactic acid production as follows: 5 mg of each PLA (all, acid-terminated and purchased from PolySciTech (W. Lafayette, USA)), P(D)LA 10-15,000 Da, P(D,L,)LA (Resomer R202H, 10-18,000 Da) or P(L)LA 15-25,000 Da) suspended in 0.5 mL of 0.4 M Tris-HCl (pH 8.0) were mixed with 50 μg of purified enzyme and incubated for 48 h at 37 ºC with shaking (1000 rpm). Samples were then centrifuged at 12,000 g for 5 min at 4 ºC. 200 μl of supernatant were mixed with 200 μl of mobile phase (0.005 N H_2_SO_4_). Sample was filtered through 13 mm Millipore PES syringe membrane filter (0.02 μm pore diameter) and analysed by HPLC Shimadzu, Prominence-I (Milton Keynes, UK) with an ion exchange column Hi PlexH (300 × 7.7 mm) (Agilent, Cheadle, UK) and 0.6 mL min^-1^ flow rate at 55 ºC (oven temperature) with UV detector set at 190-210 nm.

### Protein crystallization and structure determination

Native metagenomic esterases were purified using metal-chelate affinity chromatography, and crystallization was performed at room temperature using the sitting-drop vapor diffusion method (protein concentration 25 mg/ml, reservoir solution 0.1 M citric acid, pH 3.5 and 19% PEG 3350). The crystal was cryoprotected by transferring into paratone oil and flash frozen in liquid nitrogen. Diffraction data for the IS11 crystal was collected at 100 K at a Rigaku home source Micromax-007 with R-AXIS IV++ detector and processed using HKL3000 (48). The structure was solved by molecular replacement using Phenix.phaser (49) and a model built by AlphaFold2 (50). Model building and refinement were performed using Phenix.refine and Coot (51). TLS parameterization was utilized for refinement, and *B*-factors were refined as isotropic. Structure geometry and validation were performed using the Phenix Molprobity tools. Data collection and refinement statistics for this structure are summarized in Table S3.

### Accession numbers

SSU rRNA gene sequences were deposited to GenBank as BioProject ID: PRJNA881593. Sequences of IS10-IS12 proteins were deposited to GenBank under accession numbers OL304252, OL304253, and OL304254. The atomic coordinates of IS11 have been deposited in the Protein Data Bank (PDB), with accession code 7SPN.

## RESULTS and DISCUSSION

### Natural microbial communities of terrestrial hydrothermal vents of Ischia and effect of polyester enrichments

To provide insights into the composition of natural microbial communities and thermophilic enzymes of hydrothermal vents of the island of Ischia, four sediment samples were collected from the Cavascura hot spring (samples IS1 and IS2) and from Maronti beach near Sant’Angelo (samples IS3 and IS4) (see Materials and Methods). Both sites represent thermophilic habitats with slightly different environmental conditions: IS1 (pH 7.0, 45 ºC), IS2 (pH 8.5, 55 ºC), IS3 (pH 4.5, 75 ºC), and IS4 (pH 5.0, 85 ºC) (Table S1). From each sample, total DNA was extracted and subjected to barcoded amplicon sequencing of the V4 region of 16S rRNA gene. Sequence analysis revealed that the IS1 community comprised mainly *Pseudomonas* (17.2 %), class Anaerolineae (Chloroflexi) (12.3%), class Armatimonadota (10.0%), *Elizabethkingia* (phylum Bacteroidota) (9.5%), other Myxococcota (9.1 %), *Sphingobacterium* (order Sphingobacteriales, class Bacteroidia, phylum Bacteroidota) (6.7 %), and class Nitrospirota (6.4%), whereas the IS2 community was dominated by *Caldimonas* (order Burkholderiales, class Gammaproteobacteria) (63.9 %), *Cutibacterium* (order Propionibacteriales, class Actinobacteria) (17.2%), and *Thermus* (phylum Deinococcota) (16 %) (Figure 1). In contrast, the IS3 community was mainly represented by Bacillales (Firmicutes), namely *Brevibacillus* (48.3%) and *Geobacillus* (42%), and other Bacilli (4.4 %), whereas IS4 comprised *Sphingobacterium* (Sphingobacteriales, Bacteroidetes) (31.9 %), *Thermobaculum* (Thermobaculales, Chloroflexi) (17.4 %) and *Geobacillus* (10.7 %), followed by *Pseudomonas* (7%) and *Bacillus* (6.1%) (Figure 1). The observed differences in the taxonomic composition of the Cavascura (IS1 and IS2) and Maronti (IS3 and IS4) samples can be attributed to different environmental conditions (temperature and pH) at the sampling sites.

**Figure 1.**
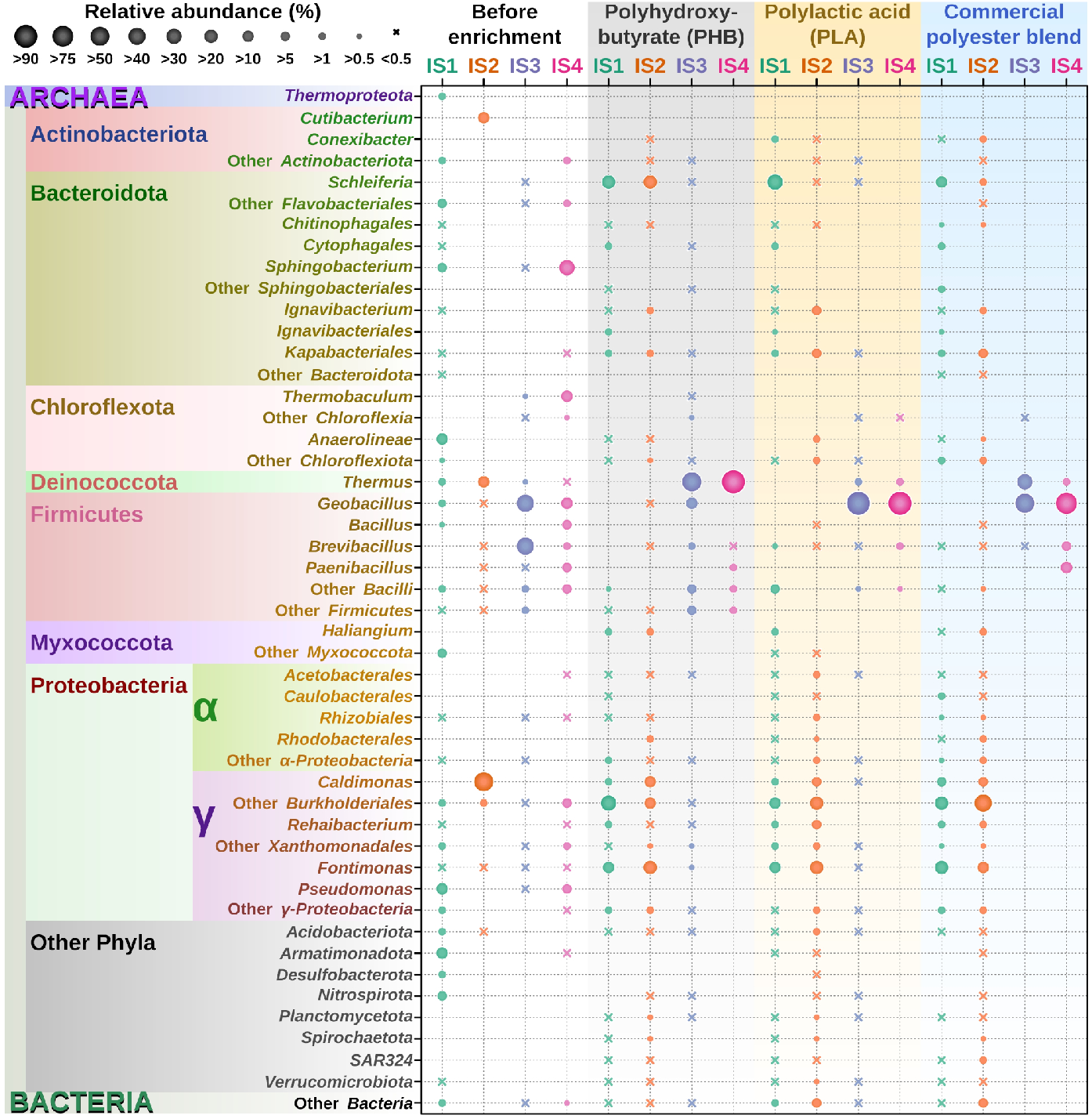
The composition of microbial communities of native samples (IS1-IS4) and polyester enrichment cultures from the Ischia hydrothermal vents. as revealed using barcoded amplicon sequencing of the V4 region of 16S rRNA gene.

Using the four sediment samples from two Ischia sites, twelve enrichment cultures were established with different polyester plastics as carbon substrate including polyhydroxybutyrate (PHB), polylactic acid (PLA) and commercial polyester blend (Table S1). After two weeks of incubation with polyesters, the IS1 enrichment culture showed a drastic increase in the relative abundance of members of the order Burkholderiales within the families Comamonadaceae and Rhodocyclaceae (relative abundance 15.2-35.1 % across the three plastic enrichments), *Fontimonas* (Solimonadaceae, 16.9-27.5%), and *Schleiferia* (order Flavobacteriales, 15.6-34.4%) (Figure 1). Likewise, IS2 enrichment showed an increase in *Fontimonas* (11-26%), *Schleiferia* (21% in PLA enrichment), whereas the relative content of the *Caldimonas* decreased from 63 % to 5.7 %, in favour of members of other families of the order Burkholderiales, namely Rhodocyclaceae, Hydrogenophilaceae and Comamonadaceae (18-43%) Kapabacteriales (phylum Bacteroidota) 2.9-8% and Rehaibacterium (order Xanthomonadales) 0.3-9.3% (Figure 1). The enrichment culture with the compostable P3 blend stimulated the growth of Rhodocyclales, as both IS1 and IS2 showed a strong increase in *Thauera* compared to experiments with PHB and PLA (Figure 1). In the enrichment cultures IS3 and IS4, higher incubation temperature (75°C) selected for thermophilic bacteria, and the nature of polyester used for enrichments influenced the microbial composition (Figure 1). The PHB enrichment stimulated growth of *Thermus* (Deinococcota), which accounted for 66.7 % (92-fold increase) and 90.9% (1,280-fold increase) of the total reads in IS3 and IS4, respectively, followed by *Geobacillus* and other members of Firmicutes. In contrast, the PLA culture favoured growth of *Geobacillus*, which reached a relative abundance of 95.8% in IS3 (2.3-fold increase) and 91.8% in IS4 (8.6-fold increase), followed by *Thermus* and *Brevibacillus*. Finally, the commercial polyester blend promoted growth of both *Geobacillus* (accounted for 68 % or 1.6-fold increase) and *Thermus* (accounted for 31.5% or 43.8-fold increase) in the IS3 enrichment, whereas the IS4 culture was dominated by Firmicutes, *Geobacillus* (81 %), *Paenibacillus* (11.9%), *Brevibacillus* (5.9 %), and *Thermus* (1.18%). As expected, the Shannon index of microbial diversity (a measure of diversity of species in a community) (Figure S1) revealed an overall tendency to decrease after incubation with polyester plastics, with the exception of IS2, which also showed low diversity in the native sample with the flattened rarefaction curve (Figure S1)

### Activity-based screening of the hydrothermal metagenome library from Ischia for carboxylesterase activity

After two weeks of incubation with polyesters, total DNA was extracted from the enrichment cultures and combined for the construction of the metagenomic fosmid library IS_Libr2. In order to identify carboxylesterases with high-temperature profiles, this library was screened for esterase activity with tributyrin as substrate (for carboxylesterases and lipases) at three temperatures: 37, 50 and 70 °C. Emulsified tributyrin gives a turbid appearance to the plates, and the presence of active metagenomic esterases or lipases is seen as a clear zone around the colony. After screening 3,456 clones from the IS_Libr2 library on tributyrin agar plates, 64 positive hits were identified with 19 positive clones observed at 37 °C, 27 clones at 50 °C, and 18 clones at 70 °C. Furthermore, eight esterase positive clones detected at 50 °C were found to be unique for this temperature, whereas one unique clone was found at 70 °C suggesting that these esterases are mostly active only at elevated temperatures. Following endonuclease digestion profiling and Sanger sequencing analysis, 14 non-redundant fosmids were selected for insert sequencing using the Illumina platform, and fosmid inserts were assembled with an average size of 39 kbp. Sequence analysis revealed 12 putative ORFs encoding predicted hydrolases (including peptidases, carboxylesterases, β-lactamases, serine proteases) homologous to proteins from Chloroflexi and metagenome assembled genome (MAG) affiliated to thermophilic Chloroflexi. From candidate proteins cloned in *E. coli*, three putative carboxylesterases (IS10, IS11, and IS12) were soluble, when expressed in *E. coli* cells, and the presence of carboxylesterase activity in purified proteins was confirmed using tributyrine agarose plates assay (Table 1) and were further selected for detailed biochemical characterisation. Amino acid sequences of IS10 (314 amino acids), IS11 (455 aa), and IS12 (318 aa) showed no presence of recognizable signal peptides suggesting that they are intracellular proteins. Both IS10 and IS12 belonged to the α/β hydrolase superfamily and had 56.8% sequence identity one to another, whereas IS11 showed no significant sequence similarity to IS10 and IS12 as was a member of the large family of β-lactamases and penicillin-binding proteins (Table 1). A blastP search of the nrNCBI database revealed that amino acid sequences of IS10 and IS12 were identical to two putative α/β hydrolases HEG24678 from uncultured Chloroflexi bacteria (GenBank accession numbers HEG24678.1 and HHR50377.1, respectively), whereas the IS11 sequence exhibited the highest identity (99.1%) to the putative “class A β-lactamase-related serine hydrolase” HDX58025.1 from uncultured Dehalococcoidia. Interestingly, the top homologous proteins of Ischia esterases were the proteins identified in metagenome from a deep-sea hydrothermal vent (black smoker) in the Mid-Atlantic Ridge (South Atlantic Ocean) (52). The comparison with previously characterised proteins showed the thermostable arylesterase, Are, from *Saccharolobus solfataricus* (UniProt ID B5BLW5, 306 aa) being the top homologue for IS10 (42 % sequence identity), whereas the metagenome-derived esterase Est8 (KP699699, PDB 4YPV, 348 aa) was the top characterised homologue for IS12 (56 % sequence identity) (53,54) (Fig. S2). The IS11 sequence was homologous to penicillin-binding proteins and β-lactamases with low sequence similarity to the CmcPBP from Actinobacteria *Amycolatopsis lactamdurans* (Q06317, 36 % identity) and esterase EstB from *Burkholderia gladioli* (Q9KX40, 32 % identity) (55,56). Domain and multiple sequence alignment confirmed the presence of conserved regions and motifs linked to esterase activity in lipolytic families previously described (Fig. S2 and S3). IS10 and IS12 contained an α/β hydrolase fold (PF07859), displaying the characteristic catalytic triad composed of Ser^155^, Asp^250^ and His^281^ and the conserved consensus motif G-x-S-x-G around the active site serine (22), clustering together with representatives of family IV (Fig. S2 and S3).

**Table 1.**
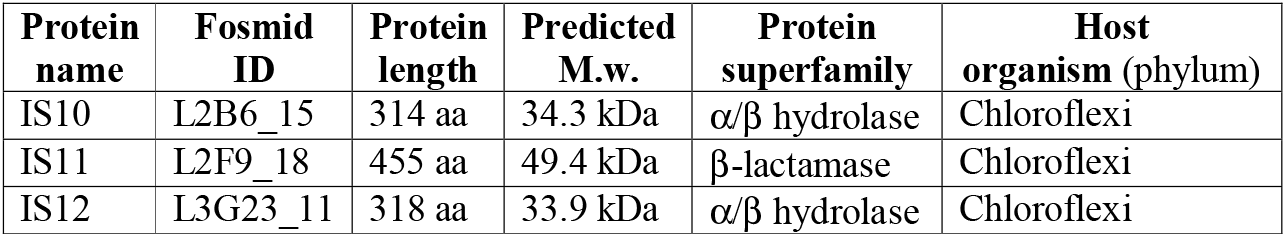
Novel carboxylesterases from the Ischia polyester enrichment metagenomes selected for biochemical and structural characterisation in this study.

The protein IS11 contained a β-lactamase domain (PF00144) and the consensus tetrapeptide S-x-x-K, perfectly conserved among all penicillin-binding enzymes and β-lactamases, surrounding the active serine Ser68. In addition, Lys71 and Tyr160 were also conserved as part of the catalytic triad of family VIII esterases, which groups enzymes with homology to class C β-lactamases and penicillin-binding proteins (Fig. S2 and S3).

### Biochemical characterisation of purified metagenomic carboxylesterases using model esterase substrates

The esterase activity of purified proteins (IS10, IS11, IS12) was initially evaluated using model esterase substrates with different chain lengths (C2-C16) at 30ºC (to diminish spontaneous substrate degradation at high temperatures). The proteins were found to be active against several short acyl chain substrates with IS10 and IS11 showing a preference to *p*-nitrophenyl butyrate (*p*NP-butyrate), α-naphthyl butyrate (αN-butyrate), and *p*NP-hexanoate, whereas IS12 was most active with *p*-NP-acetate and αN-propionate (Figure 2). All enzymes were active within a broad pH range (pH 6.0-10.0) with maximal activities at pH 9 (data not shown). The purified metagenomic carboxylesterases exhibited saturation kinetics with model esterase substrates at optimal pH (9.0) and 30 ºC (Table 2). IS10 appeared to be the most efficient esterase compared to IS11 and IS12, with the highest substrate affinity (lowest *K*_M_) and catalytic efficiency (*k*_cat_/*K*_M_) towards the tested model substrates. IS12 showed higher substrate affinity to *p*NP-butyrate and higher activity with *p*NP-acetate than IS11, whereas the latter was more active against *p*NP-butyrate (Table 2). Overall, both IS10 and IS12 appeared to be more active against the tested model substrates compared to IS11.

**Table 2.** Kinetic parameters of purified metagenomic carboxylesterases from the Ischia hydrothermal vents with model esterase substrates^a^.

| Protein | Substrate | $K_M$ (mM) | $k_{cat}$ (s<sup>-1</sup>) | $k_{cat}/K_M$ (s<sup>-1</sup> M<sup>-1</sup>) |
| --- | --- | --- | --- | --- |
| IS10 | *p*NP-acetate | 0.05 ± 0.01 | 41.97 ± 1.79 | 7.9 × 10<sup>5</sup> |
|  | *p*NP-butyrate | 0.06 ± 0.01 | 66.21 ± 3.24 | 1.2 × 10<sup>6</sup> |
|  | *p*NP-hexanoate | 0.04 ± 0.01 | 86.79 ± 3.06 | 2.0 × 10<sup>6</sup> |
| | αN-propionate | 0.06 ± 0.02 | 31.20 ± 1.93 | 5.0 × 10<sup>5</sup> |
| | αN-butyrate | 0.12 ± 0.04 | 58.60 ± 5.89 | 4.9 × 10<sup>5</sup> |
| IS11 | *p*NP-acetate | 0.53 ± 0.31 | 1.60 ± 0.30 | 3.0 × 10<sup>3</sup> |
|  | *p*NP-butyrate | 0.20 ± 0.02 | 68.81 ± 1.37 | 3.5 × 10<sup>5</sup> |
|  | *p*NP-hexanoate | 0.08 ± 0.02 | 40.28 ± 1.39 | 5.3 × 10<sup>5</sup> |
| | αN-butyrate | 0.09 ± 0.02 | 5.93 ± 0.49 | 6.9 × 10<sup>4</sup> |
| IS12 | *p*NP-acetate | 0.22 ± 0.05 | 57.10 ± 3.78 | 2.6 × 10<sup>5</sup> |
|  | *p*NP-butyrate | 0.08 ± 0.01 | 8.77 ± 0.29 | 1.1 × 10<sup>5</sup> |
|  | *p*NP-hexanoate | 0.09 ± 0.01 | 19.05 ± 0.50 | 2.1 × 10<sup>5</sup> |
| | αN-propionate | 0.69 ± 0.19 | 39.45 ± 3.7 | 5.7 × 10<sup>4</sup> |

<sup>a</sup> Reaction conditions were as indicated in Materials and Methods (pH 9.0, 30°C). Results are mean ± SD of three independent experiments. αN = α-naphthyl, *p*NP = *p*-nitrophenyl.

**Figure 2.**
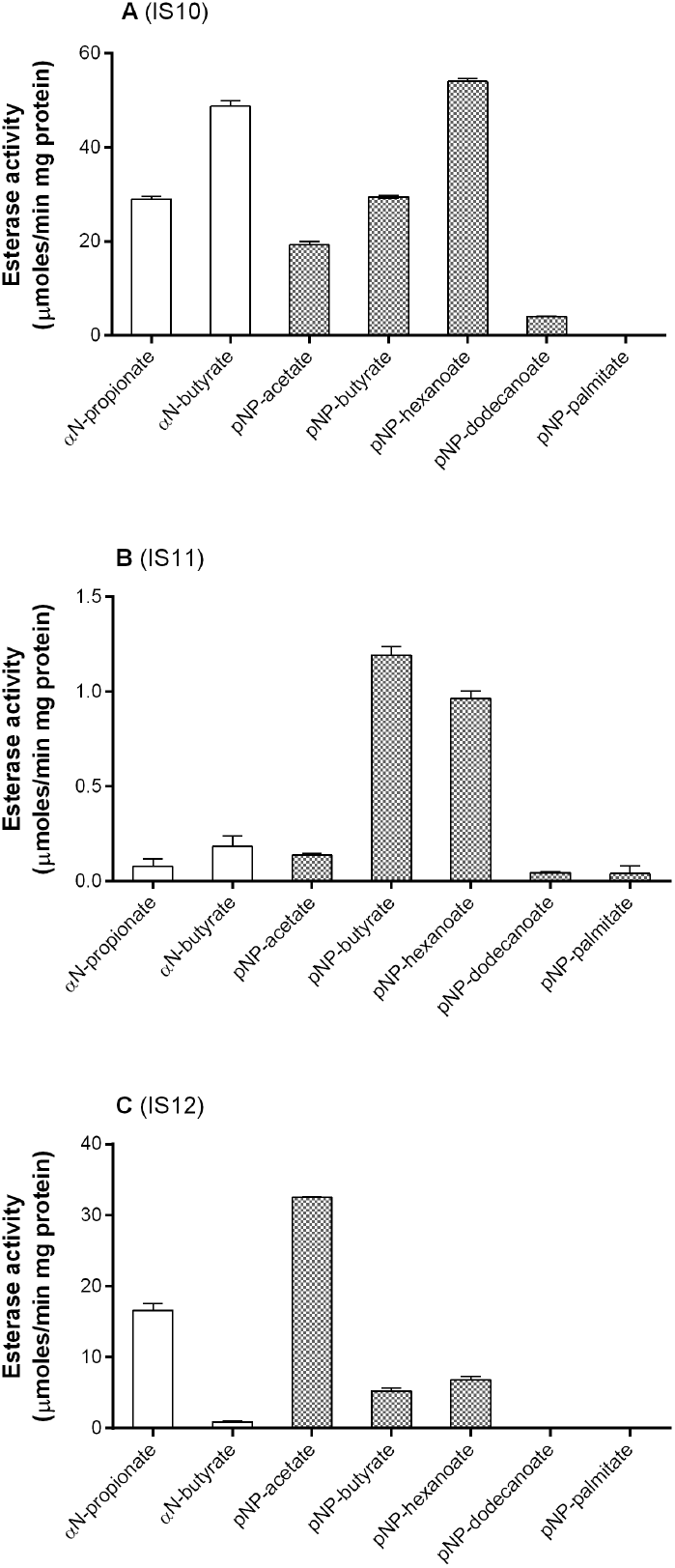
Hydrolytic activity of purified IS10 (A), IS11 (B) and IS12 (C) against model esterase substrates. The reaction mixtures contained the indicated *p*-nitrophenyl esters (*p*NP, white bars) and α-naphthyl esters (αN, grey bars) with different acyl chain lengths (reaction temperature 30ºC, see Materials and Methods for details).

Since the selected carboxylesterases originated from thermophilic environments, we investigated the effect of temperature on the activity (temperature profiles) and thermostability of purified carboxylesterases using *p*-NP-butyrate as substrate (Fig. 3). All enzymes showed significant activity at 20ºC, but reaction rates increased 5-10 times at higher temperatures with IS10 showing the highest activity at 60-70 ºC, whereas IS12 was most active at 70ºC-80ºC and IS11 at 80-90 ºC (Fig. 3). The thermostability of purified enzymes was analysed using 20 min preincubation at different temperatures (from 30 to 95ºC) followed by esterase assays with *p*NP-butyrate at 30 ºC. IS10 retained 60% activity after preincubation at 50 ºC and showed a complete loss of activity at 80ºC (Fig. 3). In contrast, both IS11 and IS12 revealed a significant decrease of activity only after 20 min preincubation at 90 ºC and 70 ºC, respectively. After two hours of incubation at 70 ºC, IS12 retained 50% of initial activity, but was completely inactivated at 80 ºC (Fig. 4). However, IS11 showed no loss or a small reduction of activity at 70 ºC and 80 ºC, respectively, and required over three hours of incubation at 90ºC for inactivation (Fig. 4). Thus, the metagenomic carboxylesterases from the Ischia hydrothermal vents are the thermophilic enzymes highly active at 70-80 ºC with IS11 and IS12 also showing significant thermostability at temperatures from 60 to 80 ºC. Furthermore, the thermostability of IS11 and IS12 was comparable with, or exceeded the, thermostability of other metagenomic esterases identified in high-temperature environments (40,57-60).

**Figure 3.**
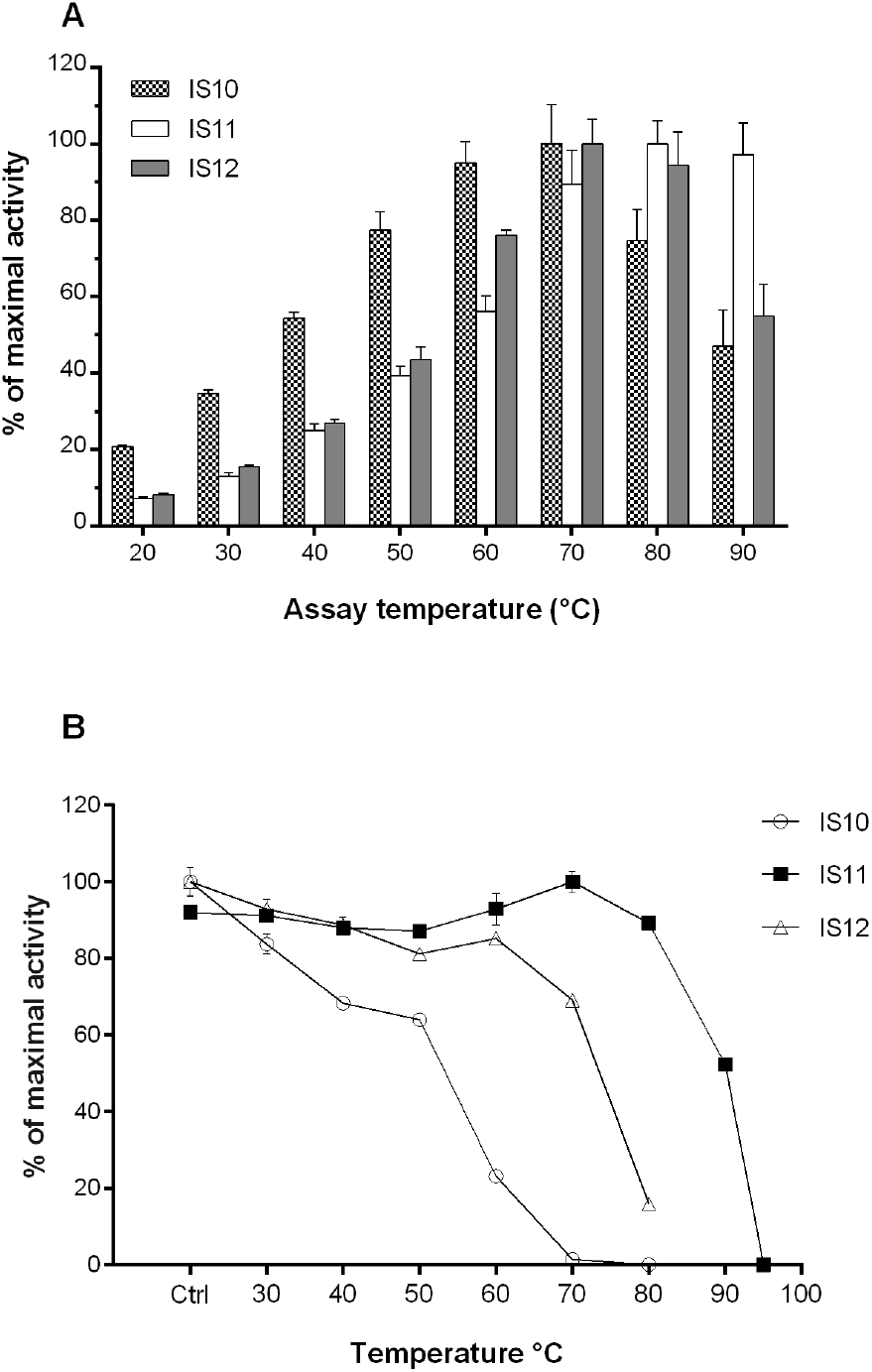
Activity temperature profiles and thermostability of purified metagenomic carboxylesterases from Ischia. **(A)** Esterase activity of purified enzymes with *p*NP-butyrate at different temperatures. **(B)** Thermostability of purified enzymes measured as residual activity after 20 min preincubation at different temperatures. Esterase activity was determined with *p*NP-butyrate as substrate at 30 ºC.

**Figure 4.**
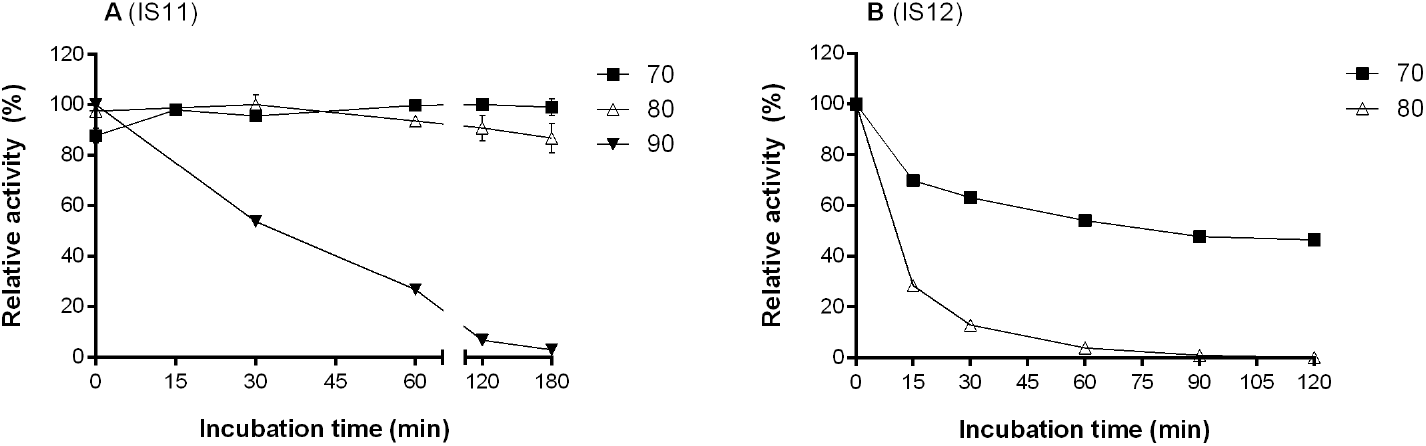
Thermoinactivation of purified IS11 (A) and IS12 (B) at different temperatures. Activity data are presented as relative activity from triplicate measurements ± SD. Residual activity was determined with *p*NP-butyrate at 30 ºC.

Esterase activity of purified metagenomic esterases was inhibited by high concentrations of NaCl (50-67% of remaining activity in the presence of 0.5 M NaCl) with IS11 showing a slightly higher resistance (Fig. S4). Similarly, IS11 retained higher activity in the presence of non-ionic detergents (43% and 53% in the presence of 2% Triton X-100 and Tween 20) (Fig. S4). With organic solvents, IS10 was inhibited by acetone, acetonitrile, ethanol, methanol, and isopropanol (10 %, v/v) (Fig. S5). In contrast, IS11 was more tolerant to these solvents (10-50 %) and was stimulated by 10% ethanol (60% increase) and 30% methanol (84% increase). Furthermore, low concentrations of these solvents (10%, v/v) stimulated esterase activity of IS12 (26-34 % increase), whereas higher concentrations of acetone and isopropanol (30%) were inhibiting. Finally, DMSO (10-30 %) stimulated esterase activity of all purified enzymes (20-46 % increase) (Fig. S5).

### Substrate range of purified carboxylesterases

To analyse the substrate range and preference of metagenomic carboxylesterases from the Ischia hydrothermal vents, the purified proteins were examined for the presence of hydrolytic activity against chemically and structurally diverse esters, including alkyl and aryl esters (Materials and Methods). Both IS10 and IS12 revealed a broad substrate range with significant activity against all 51 esters tested ester substrates and the highest activity with phenyl acetate, phenyl propionate, glyceryl tripropionate, tributyrin, and naphthyl acetate (Table S2). IS12 was also highly active toward pentane-1,5-diyl diacrylate, tri(propylene glycol) diacrylate, and vinyl propionate. The broad substrate range of IS10 and IS12 correlates with relatively large effective volumes of their active sites, 650.23 Å^3^ and 780.5 Å^3^, respectively (calculated as cavity volume/solvent accessible surface area) (23). These volumes are the largest calculated for prokaryotic esterases experimentally characterised so far, with only CalA lipase (Novozym 735) from the yeast *C. antarctica* having a larger value (23). IS11 had a more restricted substrate range, showing detectable activity against 22 ester substrates of 51 tested with a preference for benzyl (*R*)-(+)-2-hydroxy-3-phenylpropionate (Suppl. Table S2). The three metagenomic esterases revealed no apparent enantio-preference and hydrolysed both enantiomers of several tested commercially available chiral substrates.

The purified metagenomic esterases were also tested for hydrolytic activity against the T-2 mycotoxin, which contains three ester groups on its side chains. The T-2 and deacetylated HT-2 toxins are members of the large group of trichothecene mycotoxins (over 190 derivatives) containing a tetracyclic ring system (61). Mycotoxins are highly toxic fungal metabolites frequently contaminating food and feed and causing negative effects on human health, animals, and economy (62,63). While physical and chemical methods have been used to detoxify mycotoxins, biological detoxification using enzymes or microbes is more attractive due to specificity, safety, and costs. With T-2 as substrate, both IS10 and IS12 showed high esterase activity based on a pH-shift assay with phenol red (2.3 U/mg and 4.4 U/mg, respectively, at 37ºC and pH 8.0), whereas IS11 was found to be inactive. Hydrolytic activity of IS10 and IS12 against T-2 was confirmed using HPLC, which also revealed the formation of different reaction products (Fig. 5). IS10 produced HT-2 as the main product, whereas HT-2 was present as the minor product in the reaction mixture with IS12, which produced mostly the T-2 triol as the main product (Fig. 5). Since the T-2 triol is known to be less toxic than T-2 and HT-2 (64), IS12 might represent a promising candidate for the biodetoxification of T-2 and HT-2.

**Figure 5.**
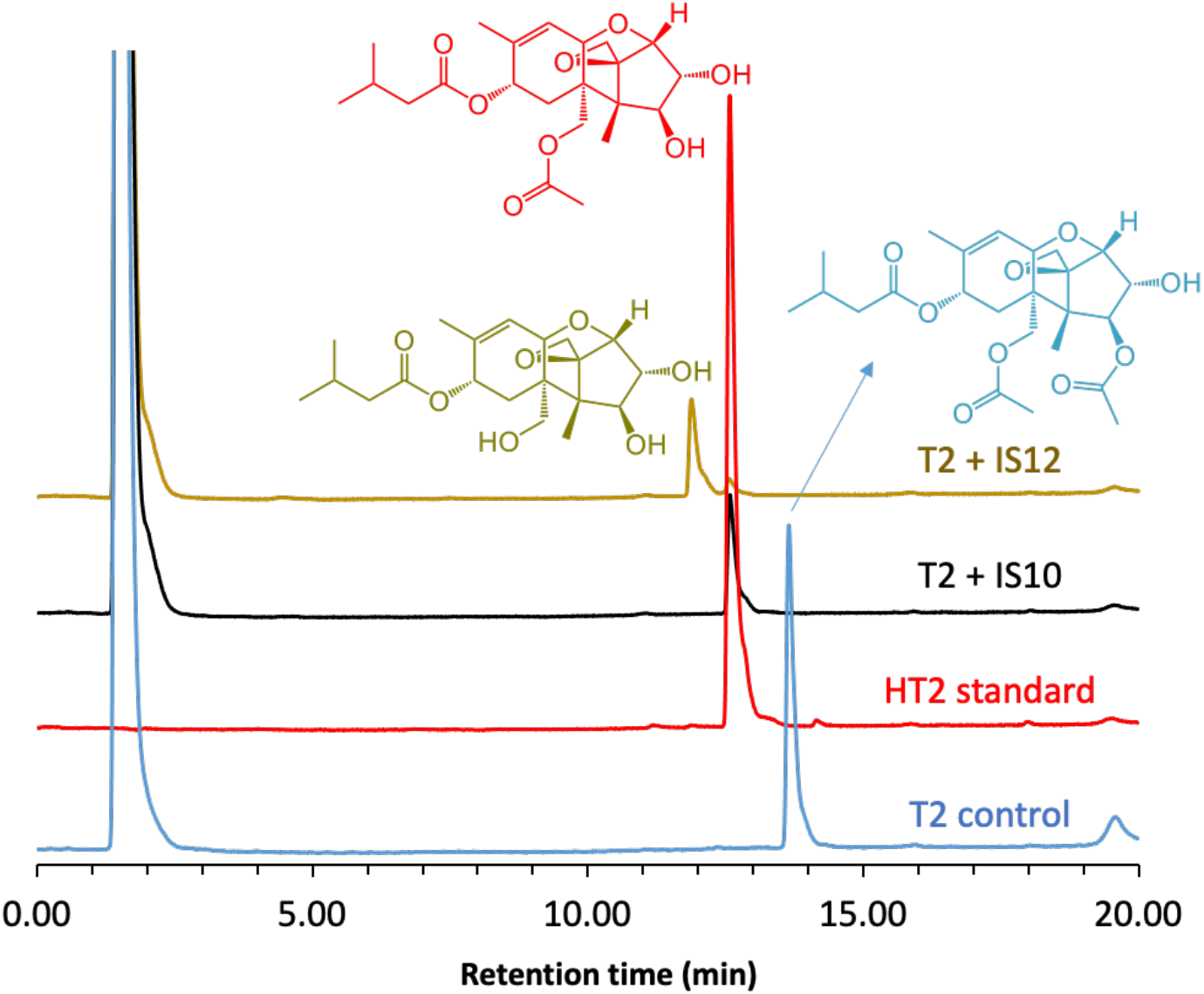
Hydrolytic activity of purified IS10 and IS12 against the mycotoxin T-2: HPLC analysis of reaction products. Purified IS10 and IS12 were incubated with T-2 (at 37 ºC and pH 8.0), and reaction products were analysed using HPLC (see Materials and Methods for experimental details).

Since our metagenomic libraries were prepared using enrichment cultures with synthetic polyesters, the purified metagenomic esterases were also tested for the presence of polyesterase activity. Although recent studies on biocatalytic depolymerization of synthetic polyesters including PLA and polyethylene terephthalate (PET) have shown the potential of microbial carboxylesterases, there is an urgent need to identify novel robust polyesterases for applications in plastics recycling (35,47,65). The purified metagenomic esterases were screened for the presence of polyesterase activity using an agarose plate assay with the emulsified PET model substrate, 3PET. These screens revealed the presence of polyesterase activity against 3PET in both IS10 and IS12, as indicated by the formation of a clear zone around the wells with loaded enzymes after incubation at 37ºC (Fig. 6A). Purified IS11 did not show a visible clearance zone on the 3PET plate, however the *in vitro* assay of hydrolysis of 3PET and HPLC analysis of reaction products, showed an increase in MHET, which was the main hydrolysis product while IS10 and IS12 produced BHET as the principal hydrolysis product. Thus, both IS10 and IS12 exhibit broad substrate profiles and were able to degrade both mycotoxins and polyesters.

**Figure 6.**
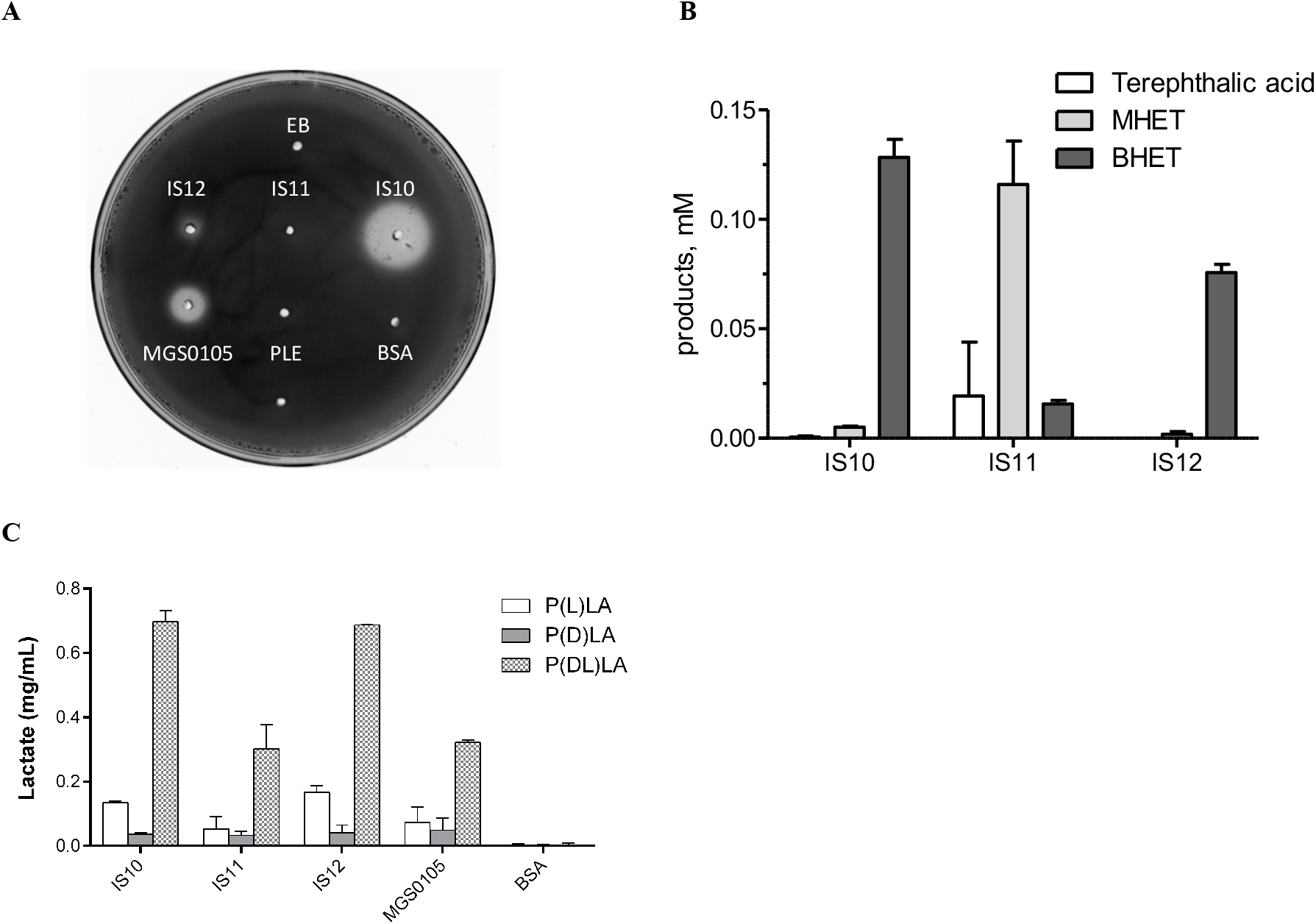
Polyesterase activity of metagenomic esterases against PLA and 3PET. **(A)** plate assay with emulsified 3PET as substrate. The formation of a clear zone around the wells with loaded enzyme indicates the presence of polyesterase activity. Agarose plates (1.5%) containing 0.2 % emulsified 3PET and loaded proteins (50 μg/well) were incubated at 37ºC and monitored for three days. Porcine liver esterase (PLE), bovine serum albumin (BSA) and elution buffer (EB) were used as a negative, MGS0105 (45) as a positive control. **(B)** HPLC assay of 3PET hydrolysis products after 16 h of incubation at 30 ºC, elution buffer was used as a negative control (not shown). **(C)** HPLC analysis of hydrolysis of PLA incubated with metagenomic esterases for 48 hrs at 30 ºC.

### Structural studies of metagenomic carboxylesterases

To provide structural insights into the active site and activity of metagenomic carboxylesterases, purified proteins (IS10, IS11, and IS12) were subjected to crystallization trials. IS11 produced diffracting crystals, and its crystal structure was determined by molecular replacement (Table S3, Materials and Methods). The overall structure of IS11 revealed a protein dimer with protomers composed of two structural domains, an N-terminal β-lactamase-like serine hydrolase domain (1-345 aa) connected via a flexible linker (346-358 aa) to a C-terminal lipocalin domain (Fig. 7). Protein oligomerization has been suggested to contribute to thermostability of several thermophilic carboxylesterases (e.g. AFEst, PestE, EstE1) (33,60,66). Accordingly, the results of size-exclusion chromatography of purified IS11, as well as IS10 and IS12, suggest that these proteins exist as dimers in solution (Figure S7).

**Figure 7.**
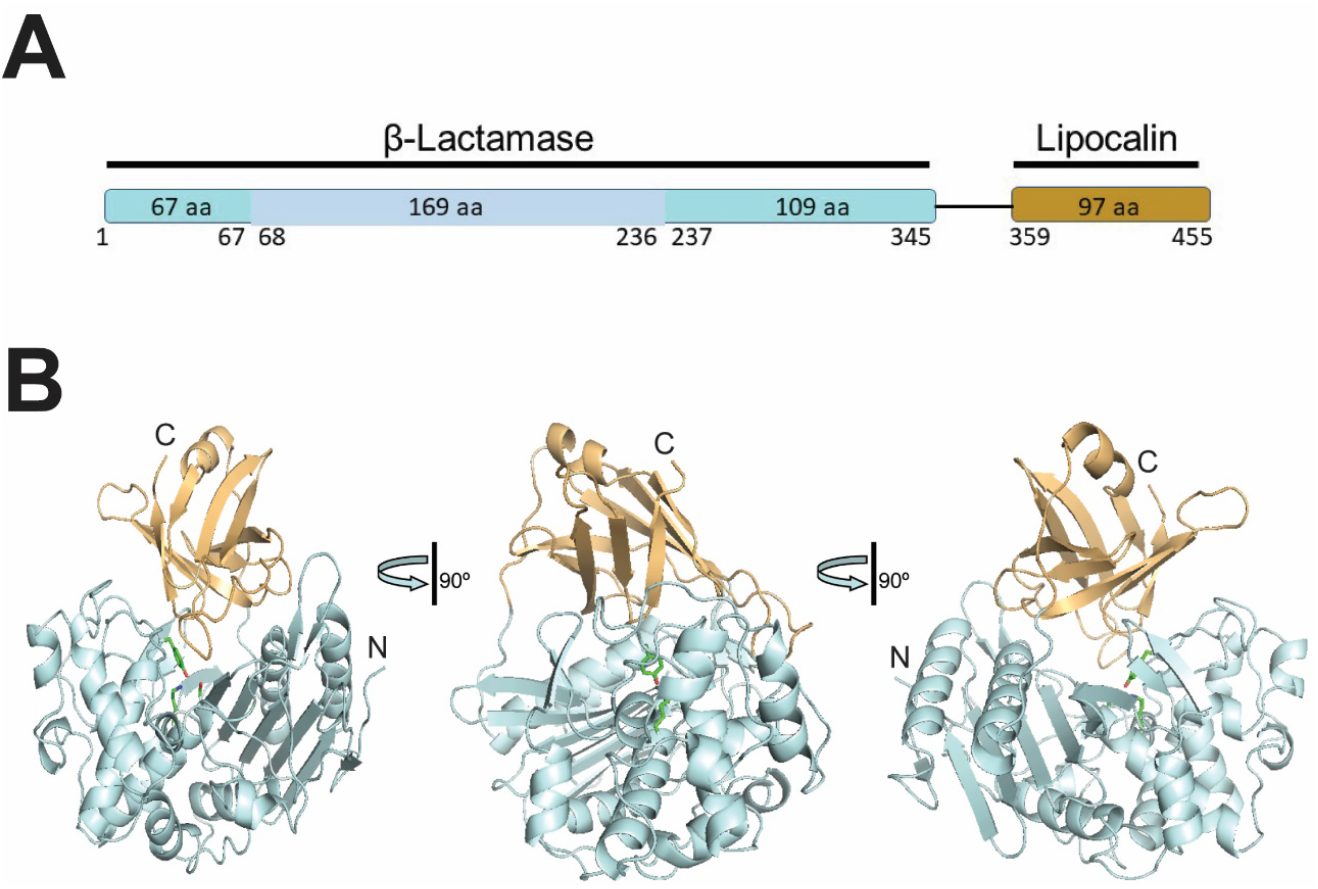
Crystal structure of IS11. (**A**), Schematic representation of the IS11 domains: the N-terminal β-lactamase related Ser hydrolase domain is colored cyan with all-helical sub-domain shown in light blue, whereas the C-terminal lipocalin domain in orange. (**B**), overall fold of the IS11 protomer shown in three views related by 90º rotations. The protein domains are shown as ribbon diagrams with the core domain (β-lactamase) colored pale cyan, whereas the C-terminal lipocalin domain is colored light orange. The position of the active site is indicated by the side chains of catalytic Ser68, Lys71, and Tyr160, whereas the protein N- and C-terminal ends are labelled (N and C).

The serine β-lactamases (classes A, C, and D) are structurally and evolutionary related to penicillin-binding proteins (the targets of β-lactam antibiotics), which also include hydrolytic DD-peptidases (67,68). The overall structure of the IS11 β-lactamase domain is composed of a mostly α-helical (all-α) sub-domain inserted into an α/β/α sandwich (or an α/β sub-domain) (Fig. 7 and 8 and Fig. S6). The α/β/α sandwich sub-domain includes a nine-stranded antiparallel β-sheet flanked by two helices on each side, whereas the mostly helical sub-domain comprises nine α-helices (Fig. 8). The two sub-domains form a groove accommodating the catalytic residues including Ser68 and Lys71 (1^st^ motif S-x-x-K), Tyr160 and Asn162 (2^nd^ motif Y-x-N/S), and His299 (3^rd^ motif H/R/K-T/S/G-G). Accordingly, the IS11 structure revealed the presence of an additional electron density positioned near the side chains of Ser68, Tyr160, and His299, represents an unknown ligand covalently attached to Ser68 (could not be modeled with various components of the protein purification or crystallization solutions) (Fig. 9a). The positioning of these catalytic residues was also conserved in the active sites of the biochemically characterised carboxylesterases with a β-lactamase fold (family VIII): EstB from *Burkholderia gladioli* and Pab87 from *Pyrococcus abyssi* (69,70), suggesting a common catalytic mechanism with acylation-deacylation. The catalytic cleft of IS11 also contains several hydrophobic and polar residues potentially involved in substrate binding (Asp126, Phe128, Trp158, Asn304, Ile307, Leu309) (Fig. 9).

**Figure 8.**
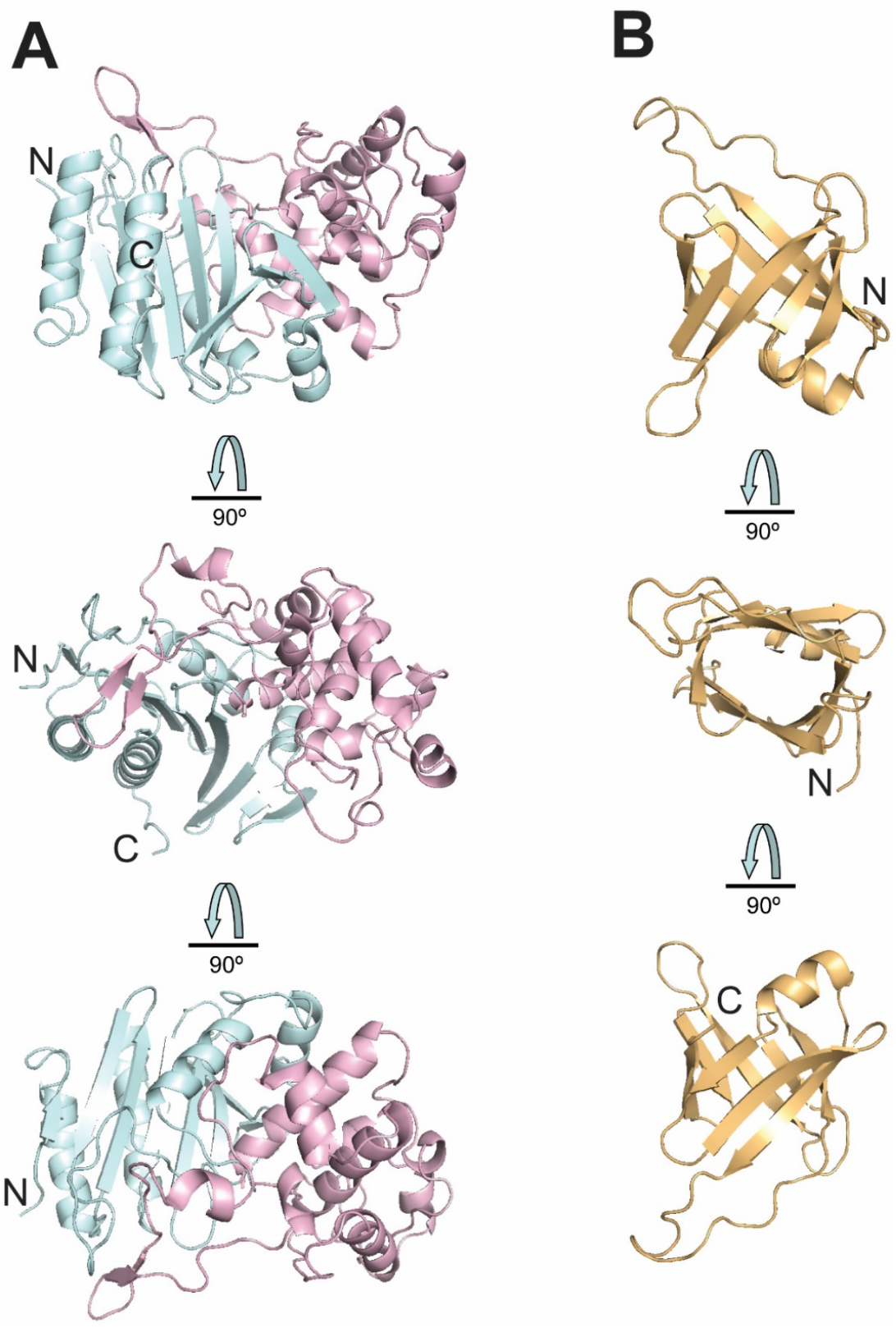
Crystal structure of the IS11 β-lactamase and lipocalin domains. (A), The N-terminal β-lactamase-like domain with two sub-domains colored pale cyan (α/β) and light pink (all-helical). (B), The lipocalin domain. The domains are shown in three views related by 90º rotations with the N- and C-termini labelled (N and C).

**Figure 9.**
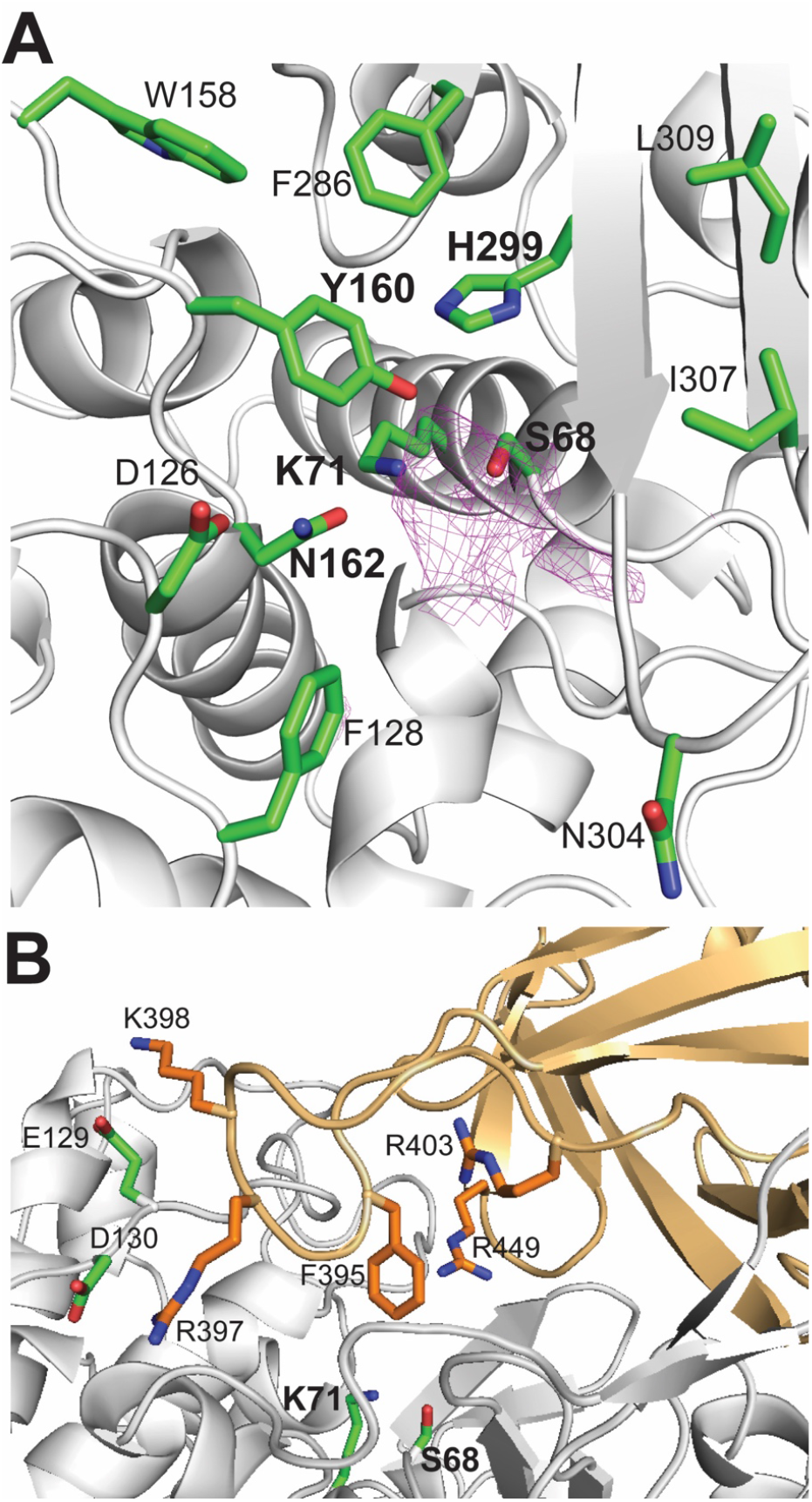
Close-up view of the IS11 active site. (A), The core domain showing the active site cleft with catalytic residues: motif-1 (Ser68 and Lys71), motif-2 (Tyr160 and Asn162), and motif-3 (His299). The magenta-coloured mesh represents an additional electron density (a 2Fo-Fc omit map contoured at 2.5 σ) covalently attached to the Ser68 side chain. (B), The proline-rich loop of the lipocalin domain covering the active site and residues potentially contributing to substrate binding. Protein ribbon diagrams are coloured grey (the β-lactamase domain) and light orange (the lipocalin domain), whereas the side chains of residues are shown as sticks with green and orange carbons, respectively

The C-terminal domain of IS11 represents a typical lipocalin fold with one α-helix and an eight-stranded antiparallel β-barrel containing a hydrophobic core (Fig. 10). Lipocalins are a diverse family of small individual proteins or domains (160-180 aa), which bind various hydrophobic molecules (e.g. fatty acids) in a binding pocket located inside the barrel (71). Although lipocalins are very divergent in their sequences and functions, their structures exhibit remarkable similarity. The lipocalin α-helix of IS11 closes off the top of the β-barrel, whose interior represents a ligand-binding site coated mostly with hydrophobic residues (Fig. 10). In the IS11 protomer, the lipocalin domain covers the β-lactamase domain shielding the catalytic cleft with the extended proline-rich strand (Pro391-Ser409) containing eight Pro residues (Pro391, Pro396, Pro401, Pro402, Pro404, Pro406, Pro407, and Pro408) (Fig. 10). In the thermophilic carboxylesterase Est2 from *Alicyclobacillus acidocaldarius*, the increased number of Pro residues has been suggested to be important for thermostability, because they reduce the flexibility of loops and other structural elements making them more resistant to denaturation (72). The side chains of several residues of the lipocalin domain and proline-rich strand are positioned close to the IS11 active site suggesting that they can be involved in substrate binding (Phe395, Arg397, Arg398, Arg403, Arg449). Typically for all lipocalins, the interior of the IS11 β-barrel is coated by mostly hydrophobic and polar residues (Leu373, Ser374, Ile376, Leu387, Gln389, Leu426, Ser429, Phe444, Phe446, Phe451). Proline-rich sequences are also known to be directly involved or facilitating protein-protein interactions or oligomerization (73). However, the IS11 dimer structure revealed no obvious interactions between the individual lipocalin domains suggesting that the lipocalin domain of IS11 participates in substrate binding, rather than in the oligomerization.

**Figure 10.**
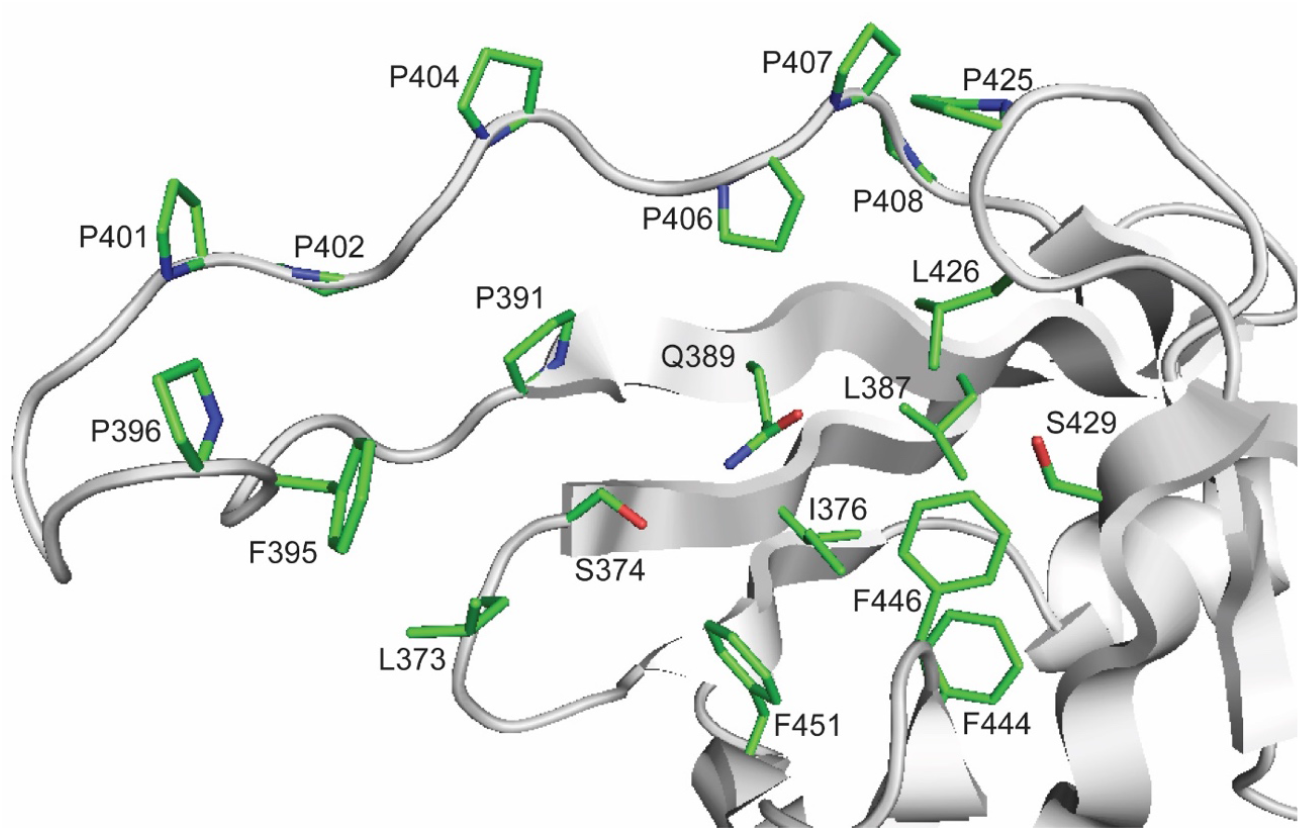
Crystal structure of the IS11 lipocalin domain: ligand binding site and proline-rich loop. The protein ribbon diagram is coloured in grey with the residues of ligand binding pocket shown as sticks with green carbons and labelled.

High-quality structural models of IS10 and IS12 proteins constructed using the Phyre2 server (Fig. S8) revealed the presence of a core domain with a classical α/β hydrolase fold and an all-helical domain, as well as a serine hydrolase catalytic triad (Ser155, His280, and Asp250 in both proteins) (Fig. S9). The putative catalytic nucleophile Ser155 is located on the classical nucleophilic elbow, a short sharp turn between a β-strand and α-helix. It is located at the bottom of the active site, which is mostly covered by the all-helical lid domain (Fig. S9). Both acyl- and alcohol-binding pockets of IS10 and IS12 include several hydrophobic and polar residues potentially involved in substrate binding (IS10: His81, Trp85, His93, Asn159, Tyr183, Val185, Leu252; IS12: Trp85, Ile87, His93, Asn159, Tyr183, Leu252, Ile279, Val283, Trp284, Leu285) (Fig. S9). Furthermore, the lid domains of both enzymes contain additional hydrophobic residues, which can contribute to substrate binding (IS10: Phe34, Met38, Phe203, Leu204, Met208, Met209, Tyr211; IS12: Phe22, Met34, Tyr195, Leu203, Leu204, Met209, Phe212, Trp213).

## CONCLUSION

Present work has demonstrated a high value of high-temperature microbial habitats, particularly of the volcanic island of Ischia (Italy), Terme di Cavascura and Maronti Beach hydrotherms populated by taxonomically diverse microorganisms, as a resource for discovery of high-temperature active enzymes. As revealed by an in-depth characterization of three metagenomics-derived carboxylesterases (IS10, IS11 and IS12) they were active at temperatures as high as 70-90 ºC and were capable to degradation of bio-based and synthetic polyester plastics. The 3PET was hydrolysed by IS10 and IS12 to predominantly bis(2-hydroxyethyl)terephthalate (BHET), while IS11 produced mono(2-hydroxyethyl)terephthalate (MHET) as a main product. Interestingly, IS12 further degraded mycotoxin T-2, a common agent causing poisoning the animal feed, to the less toxic T-2 triol. The three wild-type enzymes may readily be applicable in pilot trials in industrial processes relevant to the circular bioeconomy for plastics and/or in the production of toxin free foods and feeds. This study can also serve as a starting point for deepening our knowledge on structural determinants for substrate specificity in carboxylesterases and for rational engineering to further improve their catalytic efficiencies to make them accepting PET oligomers larger than 3PET.

## Supporting information

GenBank submission

## ACKNOWLEDGMENTS

This study was conducted under the auspices of the FuturEnzyme Project funded by the European Union’s Horizon 2020 Research and Innovation Programme under Grant Agreement No. 101000327. M.F. also acknowledges financial support under Grants PID2020-112758RB-I00 and PDC2021-121534-I00 from the Ministerio de Ciencia e Innovación, Agencia Estatal de Investigación (AEI) (Digital Object Identifier 10.13039/501100011033), Fondo Europeo de Desarrollo Regional (FEDER) and the European Union (“NextGenerationEU/PRTR”), and Grant 2020AEP061 (M.F.) from the Agencia Estatal CSIC. C. Coscolín thanks the Ministerio de Economía y Competitividad and FEDER for a PhD fellowship (Grant BES-2015-073829). P.N.G. and A.F.Y. acknowledge the Natural Environment Research Council (NERC) - funded Plastic Vectors project NE/S004548/1 and the Sêr Cymru programme partly funded by the ERDF through the Welsh Government for the support of the project BioPOL4Life.

## Notes

https://dataview.ncbi.nlm.nih.gov/object/PRJNA881593?reviewer=6amc46gk673bqn5ph2tuq5d2br

### Competing Interest Statement

The authors have declared no competing interest.

### Summary of Updates

some change in the Figure 9, its figure legend, and text discussing Fig 9a; BioProject number amended

## REFERENCES

1. Kyrpides NC, Hugenholtz P, Eisen JA, Woyke T, Goker M, Parker CT, Amann R, Beck BJ, Chain PS, Chun J, Colwell RR, Danchin A, Dawyndt P, Dedeurwaerdere T, DeLong EF, Detter JC, De Vos P, Donohue TJ, Dong XZ, Ehrlich DS, Fraser C, Gibbs R, Gilbert J, Gilna P, Glockner FO, Jansson JK, Keasling JD, Knight R, Labeda D, Lapidus A, Lee JS, Li WJ, Ma J, Markowitz V, Moore ER, Morrison M, Meyer F, Nelson KE, Ohkuma M, Ouzounis CA, Pace N, Parkhill J, Qin N, Rossello-Moř R, Sikorski J, Smith D, Sogin M, Steven R, Stingl U, Suzuki K, Taylor D, Tiedje JM, Tindall, B, Wagner M, Weinstock G, Weissenbach J, White O, Wang J, Zhang L, Zhou YG, Field D, Whitman WB, Garrity GM, Klenk HP. 2014. Genomic encyclopedia of bacteria and archaea: sequencing a myriad of type strains. PLoS Biol 12:e1001920

2. Yarza P, Yilmaz P, Pruesse E, Glockner FO, Ludwig W, Schleifer KH, Whitman WB, Euzeby J, Amann R, Rossello-Mora R. 2014. Uniting the classification of cultured and uncultured bacteria and archaea using 16S rRNA gene sequences. Nat Rev Microbiol 12:635–645

3. Rappe MS, Giovannoni SJ. 2003. The uncultured microbial majority. Annu Rev Microbiol 57:369–394

4. Torsvik V, Goksoyr J, Daae FL. 1990. High diversity in DNA of soil bacteria. Appl Environ Microbiol 56:782–787

5. Handelsman J. 2004. Metagenomics: application of genomics to uncultured microorganisms. Microbiol Mol Biol Rev 68:669–685

6. Ferrer M, Golyshina O, Beloqui A, Golyshin PN. 2007. Mining enzymes from extreme environments. Curr Opin Microbiol 10:207–214

7. Uchiyama T, Miyazaki K. 2009. Functional metagenomics for enzyme discovery: challenges to efficient screening. Curr Opin Biotechnol 20:616–622

8. Turnbaugh PJ, Gordon JI. 2008. An invitation to the marriage of metagenomics and metabolomics. Cell 134:708–713

9. Venter JC, Remington K, Heidelberg JF, Halpern AL, Rusch D, Eisen JA, Wu D, Paulsen I, Nelson KE, Nelson W, Fouts DE, Levy S, Knap AH, Lomas MW, Nealson K, White O, Peterson J, Hoffman J, Parsons R, Baden-Tillson H, Pfannkoch C, Rogers YH, Smith HO. 2004. Environmental genome shotgun sequencing of the Sargasso Sea. Science 304:66–74

10. Rusch DB, Halpern AL, Sutton G, Heidelberg KB, Williamson S, Yooseph S, Wu D, Eisen JA, Hoffman JM, Remington K, Beeson K, Tran B, Smith H, Baden-Tillson H, Stewart C, Thorpe J, Freeman J, Andrews-Pfannkoch C, Venter JE, Li K, Kravitz S, Heidelberg JF, Utterback T, Rogers YH, Falcón LI, Souza V, Bonilla-Rosso G, Eguiarte LE, Karl DM, Sathyendranath S, Platt T, Bermingham E, Gallardo V, Tamayo-Castillo G, Ferrari MR, Strausberg RL, Nealson K, Friedman R, Frazier M, Venter JC. 2007. The Sorcerer II Global Ocean Sampling expedition: northwest Atlantic through eastern tropical Pacific. PLoS Biol 5:e77

11. Yooseph S, Sutton G, Rusch DB, Halpern AL, Williamson SJ, Remington K, Eisen JA, Heidelberg KB, Manning G, Li W, Jaroszewski L, Cieplak P, Miller CS, Li H, Mashiyama ST, Joachimiak MP, van Belle C, Chandonia JM, Soergel DA, Zhai Y, Natarajan K, Lee S, Raphael BJ, Bafna V, Friedman R, Brenner SE, Godzik A, Eisenberg D, Dixon JE, Taylor SS, Strausberg RL, Frazier M, Venter JC. 2007. The Sorcerer II Global Ocean Sampling expedition: expanding the universe of protein families. PLoS Biol 5:e16

12. Qin J, Li R, Raes J, Arumugam M, Burgdorf KS, Manichanh C, Nielsen T, Pons N, Levenez F, Yamada T, Mende DR, Li J, Xu J, Li S, Li D, Cao J, Wang B, Liang H, Zheng H, Xie Y, Tap J, Lepage P, Bertalan M, Batto JM, Hansen T, Le Paslier D, Linneberg A, Nielsen HB, Pelletier E, Renault P, Sicheritz-Ponten T, Turner K, Zhu H, Yu C, Li S, Jian M, Zhou Y, Li Y, Zhang X, Li S, Qin N, Yang H, Wang J, Brunak S, Doré J, Guarner F, Kristiansen K, Pedersen O, Parkhill J, Weissenbach J; MetaHIT Consortium, Bork P, Ehrlich SD, Wang J. 2010. A human gut microbial gene catalogue established by metagenomic sequencing. Nature 464:59–65

13. Hess M, Sczyrba A, Egan R, Kim TW, Chokhawala H, Schroth G, Luo S, Clark DS, Chen F, Zhang T, Mackie RI, Pennacchio LA, Tringe SG, Visel A, Woyke T, Wang Z, Rubin EM. 2011. Metagenomic discovery of biomass-degrading genes and genomes from cow rumen. Science 331:463–467

14. Ferrer M, Martínez-Martínez M, Bargiela R, Streit WR, Golyshina OV, Golyshin PN. 2016. Estimating the success of enzyme bioprospecting through metagenomics: current status and future trends. Microb Biotechnol 9:22–34

15. Levitt M. 2009. Nature of the protein universe. Proc Natl Acad Sci U S A 106:11079–11084

16. Godzik A. 2011. Metagenomics and the protein universe. Curr Opin Struct Biol 21:398–403

17. Dinsdale EA, Edwards RA, Hall D, Angly F, Breitbart M, Brulc JM, Furlan M, Desnues C, Haynes M, Li L, McDaniel L, Moran MA, Nelson KE, Nilsson C, Olson R, Paul J, Brito BR, Ruan Y, Swan BK, Stevens R, Valentine DL, Thurber RV, Wegley L, White BA, Rohwer F. 2008. Functional metagenomic profiling of nine biomes. Nature 452:629–632

18. Rondon MR, August PR, Bettermann AD, Brady SF, Grossman TH, Liles MR, Loiacono KA, Lynch BA, MacNeil IA, Minor C, Tiong CL, Gilman M, Osburne MS, Clardy J, Handelsman J, Goodman RM. 2000. Cloning the soil metagenome: a strategy for accessing the genetic and functional diversity of uncultured microorganisms. Appl Environ Microbiol 66:2541–2547

19. Simon C, Daniel R. 2011. Metagenomic analyses: past and future trends. Appl Environ Microbiol 77:1153–1161

20. Robertson DE, Chaplin JA, DeSantis G, Podar M, Madden M, Chi E, Richardson T, Milan A, Miller M, Weiner DP, Wong K, McQuaid J, Farwell B, Preston LA, Tan X, Snead MA, Keller M, Mathur E, Kretz PL, Burk MJ, Short JM. 2004. Exploring nitrilase sequence space for enantioselective catalysis. Appl Environ Microbiol 70:2429–2436

21. Lorenz P, Eck J. 2005. Metagenomics and industrial applications. Nat Rev Microbiol 3:510–516

22. Bornscheuer UT. 2002. Microbial carboxyl esterases: classification, properties and application in biocatalysis. FEMS Microbiol Rev 26:73–81

23. Martínez-Martínez M, Coscolín C, Santiago G, Chow J, Stogios PJ, Bargiela R, Gertler C, Navarro-Fernández J, Bollinger A, Thies S, Méndez-García C, Popovic A, Brown G, Chernikova TN, García-Moyano A, Bjerga GEK, Pérez-García P, Hai T, Del Pozo MV, Stokke R, Steen IH, Cui H, Xu X, Nocek BP, Alcaide M, Distaso M, Mesa V, Peláez AI, Sánchez J, Buchholz PCF, Pleiss J, Fernández-Guerra A, Glöckner FO, Golyshina OV, Yakimov MM, Savchenko A, Jaeger KE, Yakunin AF, Streit WR, Golyshin PN, Guallar V, Ferrer M, The Inmare Consortium. 2018. Determinants and Prediction of Esterase Substrate Promiscuity Patterns. ACS Chem Biol 13:225–234

24. Arpigny JL, Jaeger KE. 1999. Bacterial lipolytic enzymes: classification and properties. Biochem J 343(Pt 1):177–183

25. Lenfant N, Hotelier T, Velluet E, Bourne Y, Marchot P, Chatonnet A. 2013. ESTHER, the database of the alpha/beta-hydrolase fold superfamily of proteins: tools to explore diversity of functions. Nucleic Acids Res 41:D423–429

26. Littlechild JA. 2017. Improving the ‘tool box’ for robust industrial enzymes. J Ind Microbiol Biotechnol 44:711–720

27. Popovic A, Hai T, Tchigvintsev A, Hajighasemi M, Nocek B, Khusnutdinova AN, Brown G, Glinos J, Flick R, Skarina T, Chernikova TN, Yim V, Brüls T, Paslier DL, Yakimov MM, Joachimiak A, Ferrer M, Golyshina OV, Savchenko A, Golyshin PN, Yakunin AF. 2017. Activity screening of environmental metagenomic libraries reveals novel carboxylesterase families. Sci Rep 7:44103

28. Pellis A, Cantone S, Ebert C, Gardossi L. 2018. Evolving biocatalysis to meet bioeconomy challenges and opportunities. N Biotechnol 40:154–169

29. Antranikian G, Streit WR. 2022. Microorganisms harbor keys to a circular bioeconomy making them useful tools in fighting plastic pollution and rising CO2 levels. Extremophiles 26:10

30. Kruger A, Schafers C, Schroder C, Antranikian G. 2018. Towards a sustainable biobased industry - Highlighting the impact of extremophiles. N Biotechnol 40:144–153

31. Atomi H. 2005. Recent progress towards the application of hyperthermophiles and their enzymes. Curr Opin Chem Biol 9:166–173

32. Littlechild JA. 2015. Archaeal Enzymes and Applications in Industrial Biocatalysts. Archaea 2015:147671

33. Vieille C, Zeikus GJ. 2001. Hyperthermophilic enzymes: sources, uses, and molecular mechanisms for thermostability. Microbiol Mol Biol Rev 65:1–43

34. Alcaide M, Stogios PJ, Lafraya Á, Tchigvintsev A, Flick R, Bargiela R, Chernikova TN, Reva ON, Hai T, Leggewie CC, Katzke N, La Cono V, Matesanz R, Jebbar M, Jaeger KE, Yakimov MM, Yakunin AF, Golyshin PN, Golyshina OV, Savchenko A, Ferrer M; MAMBA Consortium. 2015. Pressure adaptation is linked to thermal adaptation in salt-saturated marine habitats. Environ Microbiol 17:332–345

35. Wei R, von Haugwitz G, Pfaff L, Mican J, Badenhorst CPS, Liu W, Weber G, Austin HP, Bednar D, Damborsky J, Bornscheuer UT. 2022. Mechanism-based design of efficient PET hydrolases. ACS Catalysis 12:3382–3396

36. Rizzo C, Arcadi E, Calogero R, Sciutteri V, Consoli P, Esposito V, Canese S, Andaloro F, Romeo T. 2022. Ecological and biotechnological relevance of Mediterranean hydrothermal vent systems. Minerals 12:251

37. Fadrosh DW, Ma B, Gajer P, Sengamalay N, Ott S, Brotman RM, Ravel J. 2014. An improved dual-indexing approach for multiplexed 16S rRNA gene sequencing on the Illumina MiSeq platform. Microbiome 2:6

38. Distaso MA, Bargiela R, Brailsford FL, Williams GB, Wright S, Lunev EA, Toshchakov SV, Yakimov MM, Jones DL, Golyshin PN, Golyshina OV. 2020 High representation of archaea across all depths in oxic and low-pH sediment layers underlying an acidic stream. Front Microbiol 11:2871.

39. R Core Team. 2020. R: A language and environment for statistical computing. R Foundation for Statistical Computing, Vienna, Austria. www.R-project.org/.

40. Placido A, Hai T, Ferrer M, Chernikova TN, Distaso M, Armstrong D, Yakunin AF, Toshchakov SV, Yakimov MM, Kublanov IV, Golyshina OV, Pesole G, Ceci LR, Golyshin PN. 2015. Diversity of hydrolases from hydrothermal vent sediments of the Levante Bay, Vulcano Island Aeolian archipelago identified by activity-based metagenomics and biochemical characterization of new esterases and an arabinopyranosidase. Appl Microbiol Biotechnol 99:10031–10046

41. Zhu W, Lomsadze A, Borodovsky M. 2010. Ab initio gene identification in metagenomic sequences. Nucleic Acids Res 38:e132

42. Altschul SF, Madden TL, Schaffer AA, Zhang J, Zhang Z, Miller W, Lipman DJ. 1997. Gapped BLAST and PSI-BLAST: a new generation of protein database search programs. Nucleic Acids Res 25:3389–3402

43. Edgar RC. 2004. MUSCLE: multiple sequence alignment with high accuracy and high throughput. Nucleic Acids Res 32:1792–1797

44. Kumar S, Stecher G, Li M, Knyaz C, Tamura K. 2018. MEGA X: Molecular Evolutionary Genetics Analysis across Computing Platforms. Mol Biol Evol 35:1547–1549

45. Tchigvintsev A, Tran H, Popovic A, Kovacic F, Brown G, Flick R, Hajighasemi M, Egorova O, Somody JC, Tchigvintsev D, Khusnutdinova A, Chernikova TN, Golyshina OV, Yakimov MM, Savchenko A, Golyshin PN, Jaeger KE, Yakunin AF. 2015. The environment shapes microbial enzymes: five cold-active and salt-resistant carboxylesterases from marine metagenomes. Appl Microbiol Biotechnol 99:2165–2178

46. Guinta CI, Cea-Rama I, Alonso S, Briand ML, Bargiela R, Coscolin C, Corvini P, Ferrer M, Sanz-Aparicio J, Shahgaldian P. 2020. Tuning the properties of natural promiscuous enzymes by engineering their nano-environment. ACS Nano 14:17652–17664

47. Hajighasemi M, Tchigvintsev A, Nocek B, Flick R, Popovic A, Hai T, Khusnutdinova AN, Brown G, Xu X, Cui H, Anstett J, Chernikova TN, Brüls T, Le Paslier D, Yakimov MM, Joachimiak A, Golyshina OV, Savchenko A, Golyshin PN, Edwards EA, Yakunin AF. 2018. Screening and Characterization of Novel Polyesterases from Environmental Metagenomes with High Hydrolytic Activity against Synthetic Polyesters. Environ Sci Technol 52:12388–12401

48. Minor W, Cymborowski M, Otwinowski Z, Chruszcz M. 2006. HKL-3000: the integration of data reduction and structure solution--from diffraction images to an initial model in minutes. Acta Crystallogr D Biol Crystallogr 62:859–866

49. Liebschner D, Afonine PV, Baker ML, Bunkóczi G, Chen VB, Croll TI, Hintze B, Hung LW, Jain S, McCoy AJ, Moriarty NW, Oeffner RD, Poon BK, Prisant MG, Read RJ, Richardson JS, Richardson DC, Sammito MD, Sobolev OV, Stockwell DH, Terwilliger TC, Urzhumtsev AG, Videau LL, Williams CJ, Adams PD. 2019 Macromolecular structure determination using X-rays, neutrons and electrons: recent developments in Phenix. Acta Crystallogr D Struct Biol 75:861–877

50. Jumper J, Evans R, Pritzel A, Green T, Figurnov M, Ronneberger O, Tunyasuvunakool K, Bates R, Žídek A, Potapenko A, Bridgland A, Meyer C, Kohl SAA, Ballard AJ, Cowie A, Romera-Paredes B, Nikolov S, Jain R, Adler J, Back T, Petersen S, Reiman D, Clancy E, Zielinski M, Steinegger M, Pacholska M, Berghammer T, Bodenstein S, Silver D, Vinyals O, Senior AW, Kavukcuoglu K, Kohli P, Hassabis D. 2021 Highly accurate protein structure prediction with AlphaFold. Nature 596:583–589

51. Emsley P, Cowtan K. 2004. Coot: model-building tools for molecular graphics. Acta Crystallogr D Biol Crystallogr 60:2126–2132

52. Zhou Z, Liu Y, Xu W, Pan J, Luo ZH, Li M. 2020. Genome- and Community-Level Interaction Insights into Carbon Utilization and Element Cycling Functions of Hydrothermarchaeota in Hydrothermal Sediment. mSystems 5:e00795–19

53. Park YJ, Yoon SJ, Lee HB. 2008. A novel thermostable arylesterase from the archaeon Sulfolobus solfataricus P1: purification, characterization, and expression. J Bacteriol 190:8086–8095

54. Pereira MR, Maester TC, Mercaldi GF, de Macedo Lemos EG, Hyvonen M, Balan A. 2017. From a metagenomic source to a high-resolution structure of a novel alkaline esterase. Appl Microbiol Biotechnol 101:4935–4949

55. Coque JJ, Liras P, Martin JF. 1993. Genes for a beta-lactamase, a penicillin-binding protein and a transmembrane protein are clustered with the cephamycin biosynthetic genes in Nocardia lactamdurans. EMBO J 12:631–639

56. Petersen EI, Valinger G, Solkner B, Stubenrauch G, Schwab H. 2001. A novel esterase from Burkholderia gladioli which shows high deacetylation activity on cephalosporins is related to beta-lactamases and DD-peptidases. J Biotechnol 89:11–25

57. Lewin A, Strand T, Haugen T, Klinkenberg G, Kotlar H, Valla S., Drablos F, Wentze, A. 2016. Discovery and characterization of a thermostable esterase from an oil reservoir metagenome. Adv Enzyme Res 4:68–86

58. Leis B, Angelov A, Mientus M, Li H, Pham VT, Lauinger B, Bongen P, Pietruszka J, Goncalves LG, Santos H, Liebl W. 2015. Identification of novel esterase-active enzymes from hot environments by use of the host bacterium Thermus thermophilus. Front Microbiol 6:275

59. Miguel-Ruano V, Rivera I, Rajkovic J, Knapik K, Torrado A, Otero JM, Beneventi E, Becerra M, Sánchez-Costa M, Hidalgo A, Berenguer J, González-Siso MI, Cruces J, Rúa ML, Hermoso JA. 2021. Biochemical and Structural Characterization of a novel thermophilic esterase EstD11 provide catalytic insights for the HSL family. Comput Struct Biotechnol J 19:1214–1232

60. Sayer C, Szabo Z, Isupov MN, Ingham C, Littlechild JA. 2015. The Structure of a Novel Thermophilic Esterase from the Planctomycetes Species, Thermogutta terrifontis Reveals an Open Active Site Due to a Minimal ‘Cap’ Domain. Front Microbiol 6:1294

61. Loi M, Fanell, F, Liuzzi VC, Logrieco AF, Mule G. 2017. Mycotoxin Biotransformation by Native and Commercial Enzymes: Present and Future Perspectives. Toxins Basel 9:111

62. Liu L, Xie M, We, D. 2022. Biological Detoxification of Mycotoxins: Current Status and Future Advances. Int J Mol Sci 23:1064

63. Lyagin I, and Efremenko E. 2019. Enzymes for Detoxification of Various Mycotoxins: Origins and Mechanisms of Catalytic Action. Molecules 24:2362

64. McCormick SP, Price NP, Kurtzman CP 2012. Glucosylation and other biotransformations of T-2 toxin by yeasts of the trichomonascus clade. Appl Environ Microbiol 78:8694–8702

65. Tournier V, Topham CM, Gilles A, David B, Folgoas C, Moya-Leclair E, Kamionka E, Desrousseaux ML, Texier H, Gavalda S, Cot M, Guémard E, Dalibey M, Nomme J, Cioci G, Barbe S, Chateau M, André I, Duquesne S, Marty A. 2020. An engineered PET depolymerase to break down and recycle plastic bottles. Nature 580:216–219.

66. Palm GJ, Fernández-Álvaro E, Bogdanović X, Bartsch S, Sczodrok J, Singh RK, Böttcher D, Atomi H, Bornscheuer UT, Hinrichs W. 2011. The crystal structure of an esterase from the hyperthermophilic microorganism Pyrobaculum calidifontis VA1 explains its enantioselectivity. Appl Microbiol Biotechnol 91:1061–1072

67. Sauvage E, Kerff F, Terrak M, Ayala JA, Charlier P. 2008. The penicillin-binding proteins: structure and role in peptidoglycan biosynthesis. FEMS Microbiol Rev 32:234–258

68. Lee D, Das S, Dawson NL, Dobrijevic D, Ward J, Orengo C. 2016. Novel Computational Protocols for Functionally Classifying and Characterising Serine Beta-Lactamases. PLoS Comput Biol 12:e1004926

69. Wagner UG, Petersen EI, Schwab H, Kratky C. 2002. EstB from Burkholderia gladioli: a novel esterase with a beta-lactamase fold reveals steric factors to discriminate between esterolytic and beta-lactam cleaving activity. Protein Sci 11:467–478

70. Delfosse V, Girard E, Birck C, Delmarcelle M, Delarue M, Poch O, Schultz O, Mayer C. 2009. Structure of the archaeal pab87 peptidase reveals a novel self-compartmentalizing protease family. PLoS One 4:e4712

71. Flower DR, North AC, Sansom CE. 2000. The lipocalin protein family: structural and sequence overview. Biochim Biophys Acta 1482:9–24

72. De Simone G, Galdiero S, Manco G, Lang D, Rossi M, Pedone C. 2000. A snapshot of a transition state analogue of a novel thermophilic esterase belonging to the subfamily of mammalian hormone-sensitive lipase. J Mol Biol 303:761–771

73. Kay BK, Williamson MP, Sudol M. 2000. The importance of being proline: the interaction of proline-rich motifs in signaling proteins with their cognate domains. FASEB J 14:231–241

