## Supplementary material for "Thermophilic carboxylesterases from hydrothermal vents of the volcanic island of Ischia active on synthetic and biobased polymers and mycotoxins": Suppl_file_Ischia_paper_4 12-09-2022.docx

**Supplementary Figures**

**Figure S1.** The Shannon diversity index of the native and polyester enrichment cultures of the hydrothermal microbial communities of the volcanic island of Ischia.

**Figure S2.** Multiple sequence alignment of metagenomic carboxylesterases from Ischia with characterized homologous enzymes

**Figure S3.** Phylogenetic analysis of the metagenomic carboxylesterases from Ischia: **(A)**, IS10 and IS12; **(B)**, IS11

**Figure S4.** Carboxylesterase activity of IS10, IS11, and IS12: effect of pH, NaCl, and detergents.

**Figure S5.** Effect of organic solvents on the activity of IS10, IS11 and IS12.

**Figure S6.** Overall crystal structure of IS11: three views of the IS11 dimer related by 90º rotations.

**Figure S7. Figure S7.** Oligomeric state of purified Ischia carboxylesterases.

**Figure S8.** Structural models of IS10 (A) and IS12 (B) shown in three views related by a 90º rotation.

**Figure S9.** Structural models of IS10 and IS12: close-up view of the active sites.

**Supplementary Tables**

**Table S1.** Environmental (*in situ*) and experimental conditions for metagenome samples and enrichment cultures.

**Table S2.** Esterase activities of purified Ischia carboxylesterases against various monoesters analyzed using a pH-shift assay.

**Table S3.** X-ray crystallographic statistics for the structure of IS11.

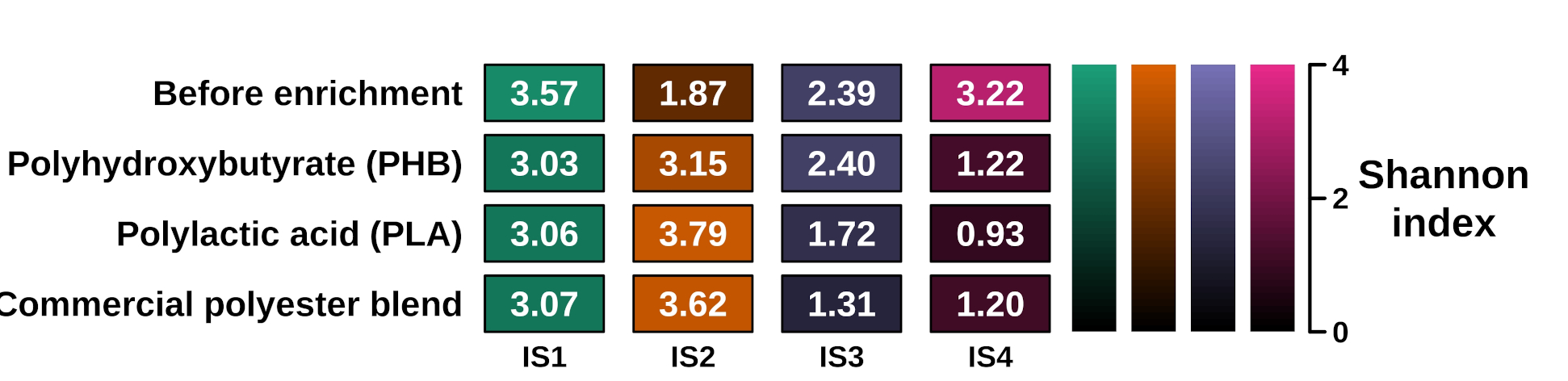

**Figure S1.** The Shannon diversity index of the native and polyester enrichment cultures of the hydrothermal microbial communities of the volcanic island of Ischia.

A

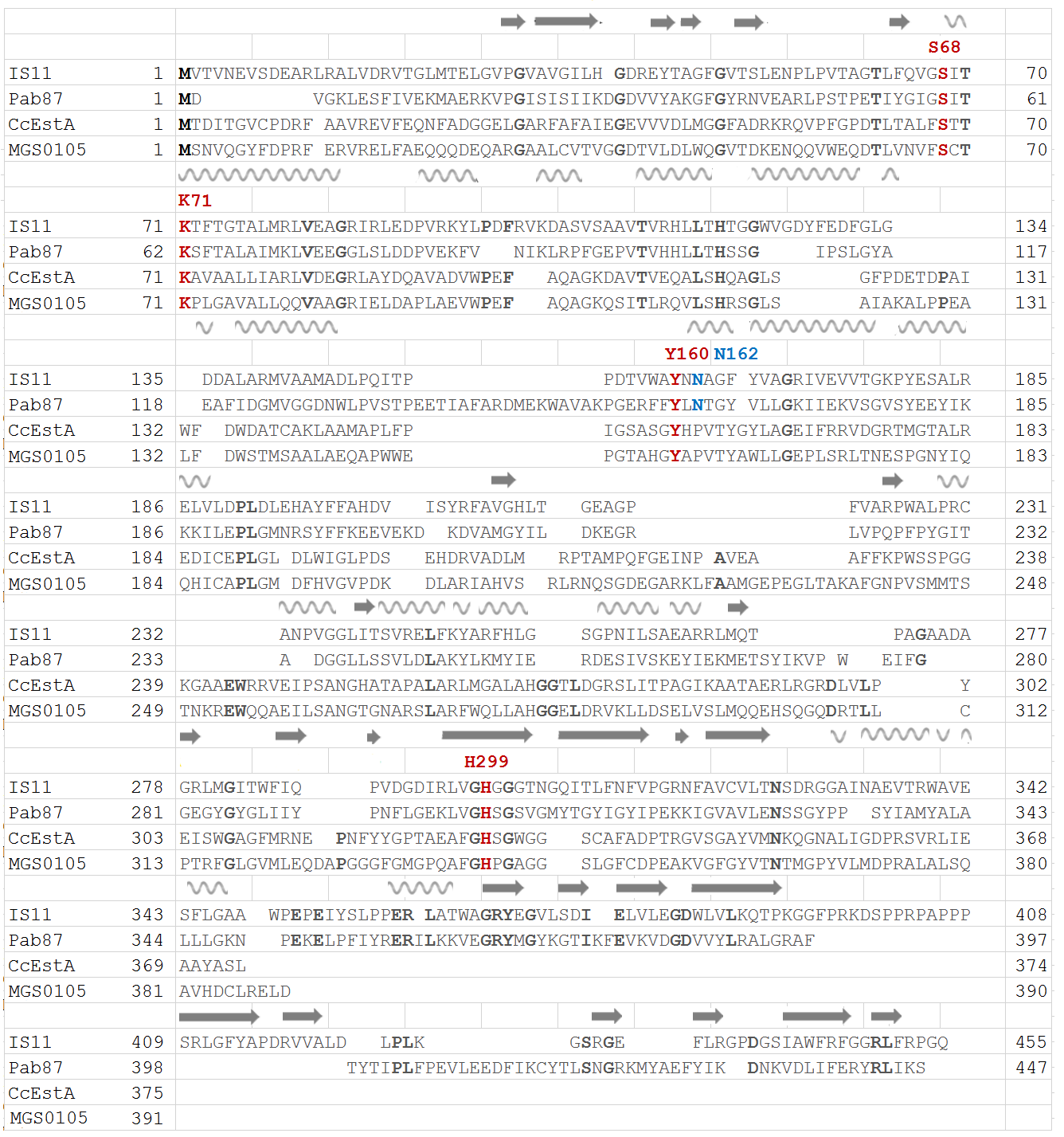

B

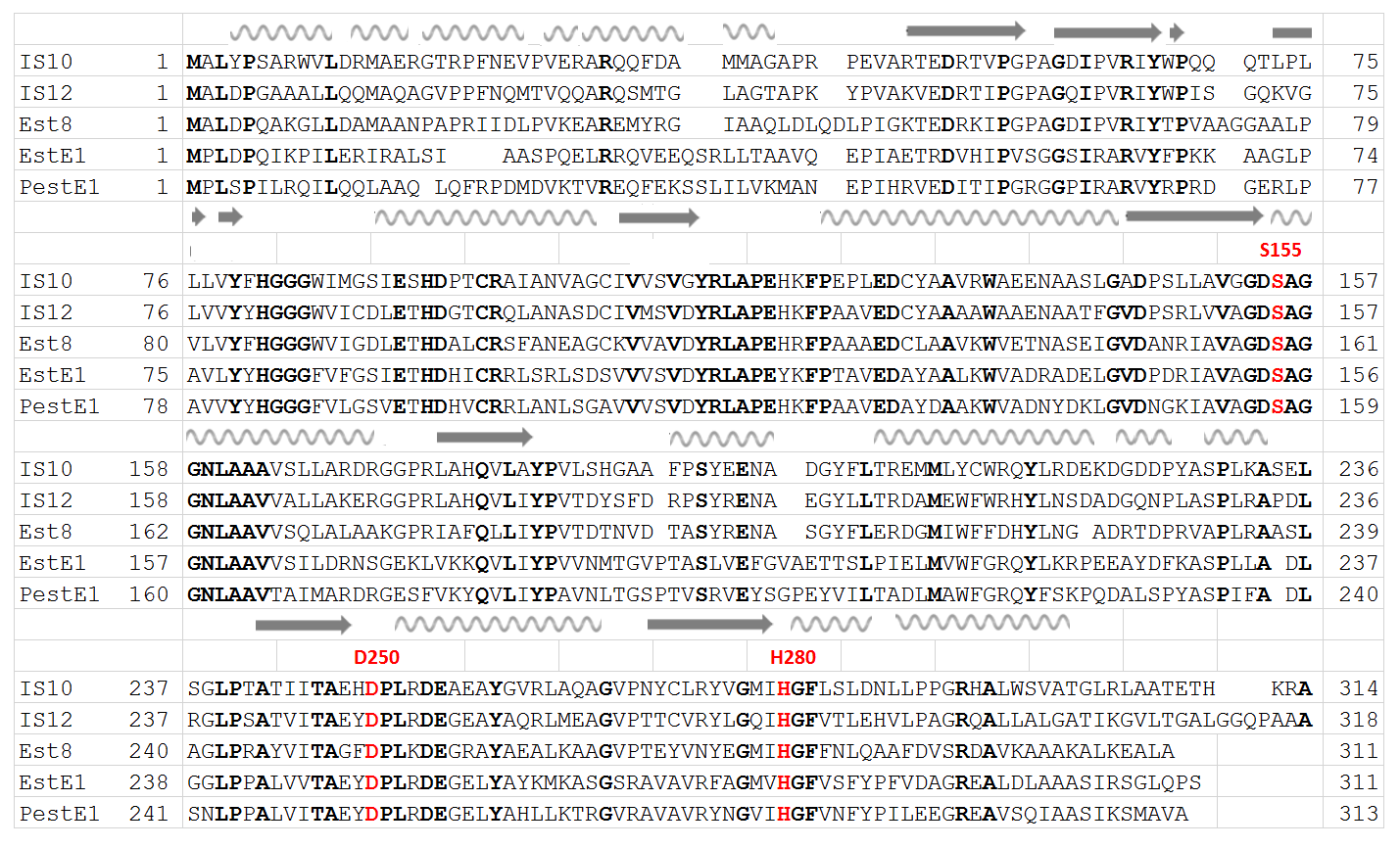

**Figure S2.** Multiple sequence alignment of metagenomic carboxylesterases from Ischia with characterized homologous enzymes. (A), IS11; (B), IS10 and IS12. The following proteins are shown: (A), IS11, Pab87 from *Pyrococcus abyssi* (Q9V2D6), MGS0105 from a cold marine metagenome (T1W153), and CcEstA from *Caulobacter crescentus* (Q9ABH2). The IS11 catalytic residues are coloured red and labelled, whereas conserved residues are shown in bold. The IS11 secondary structure elements are shown above the alignment. (B), IS10, IS12, Est8 from an oil-degrading metagenome (ALO81572), EstE1 from a thermophilic metagenome (Q5G935), and PestE1 from *Pyrobaculum calidifontis* (Q8NKS0). The catalytic residues of IS10 are coloured red and labelled, whereas the conserved residues are shown in bold. The secondary structure elements of IS11 (A) and IS10 (B) are shown above the alignments. Both sequence alignments were prepared using the MAFFT online tool (<https://mafft.cbrc.jp/alignment/server/> ) and STRAP (<http://www.bioinformatics.org/strap/> ).

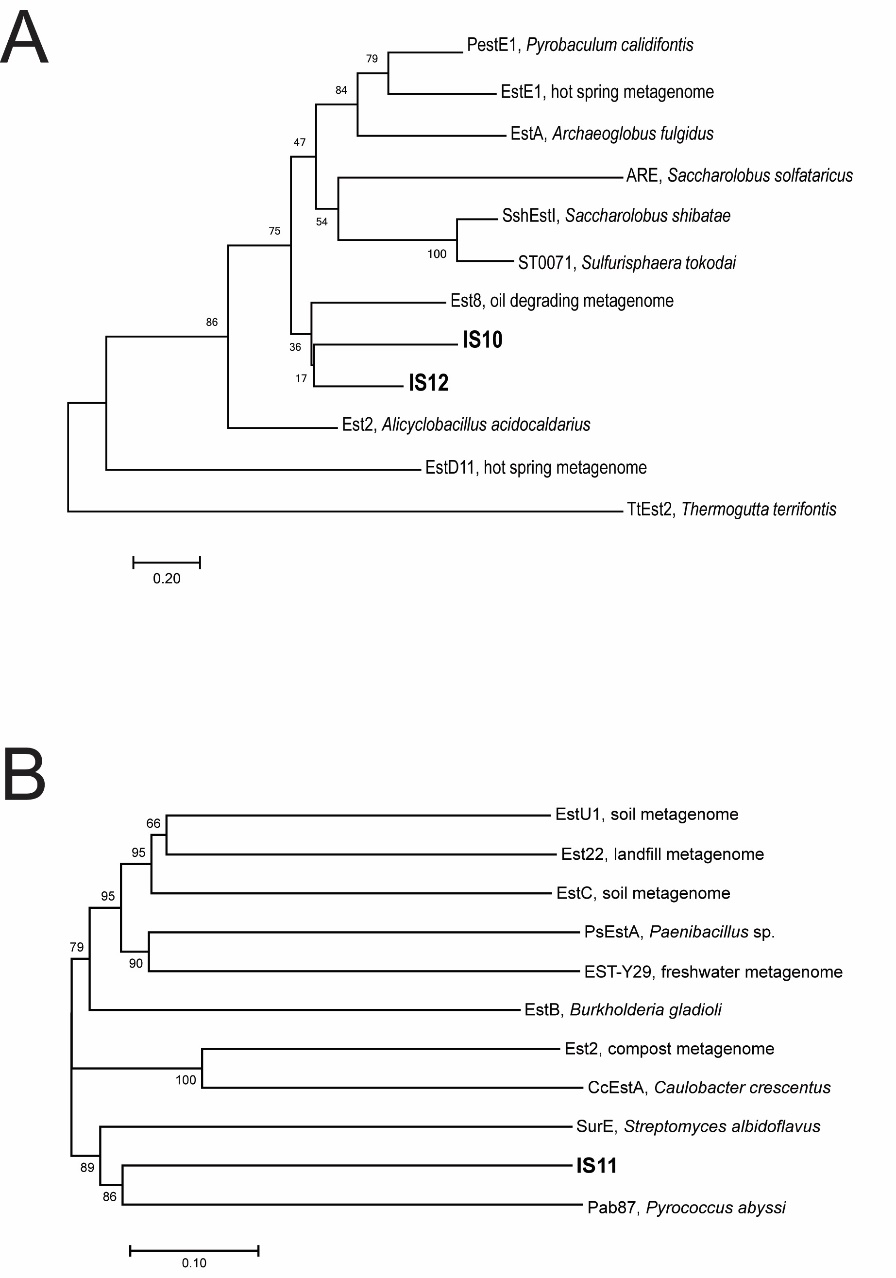

**Figure S3.** Phylogenetic analysis of the metagenomic carboxylesterases from Ischia: **(A)**, IS10 and IS12; **(B)**, IS11. The phylogenetic tree showing the relationship between IS10, IS11, IS12, and biochemically characterized carboxylesterases from various metagenomes and microorganisms. The tree was generated with a MEGA7 software using the Neighbour-Joining method (see Materials and Methods for details). The bootstraps values (for 1,000 pseudoreplicates) are indicated at the branches, and scale bars indicate 0.2 and 0.1 substitutions per position in (A) and (B), respectively. The following proteins were used for the analysis: (A), PestE1 from *Pyrobaculum calidifontis* (Q8NKS0), EstE1 from a hot spring metagenome (Q5G935), EstA from *Archaeoglobus fulgidus* (O28558), ARE from *Saccharolobus solfataricus* (B5BLW5), SshEstI from *Saccharolobus shibatae* (Q5NU42), ST0071 from *Sulfurisphaera tokodai* (Q976W8), Est8 from an oil degrading metagenome (ALO81572), Est2 from *Alicyclobacillus acidocaldarius* (Q7SIG1), EstD11 from a hot spring metagenome (A0A8I3AZT4), TtEst2 from *Thermogutta terrifontis* (KT724966); (B), EstU1 from a soil metagenome (AFU54388), Est22 from a landfill metagenome (KF052088), EstC from a soil metagenome (ACH88047), PsEstA from *Paenibacillus* sp. (AHL66978), EST-Y29 from a freshwater metagenome (NBR33940), EstB from *Burkholderia gladioli* (Q9KX40), Est2 from a compost metagenome (ALH24853), CcEstA from *Caulobacter crescentus* (Q9ABH2), SurE from *Streptomyces albidoflavus* (A0A679G4U8), Pab87 from *Pyrococcus abyssi* (Q9V2D6).

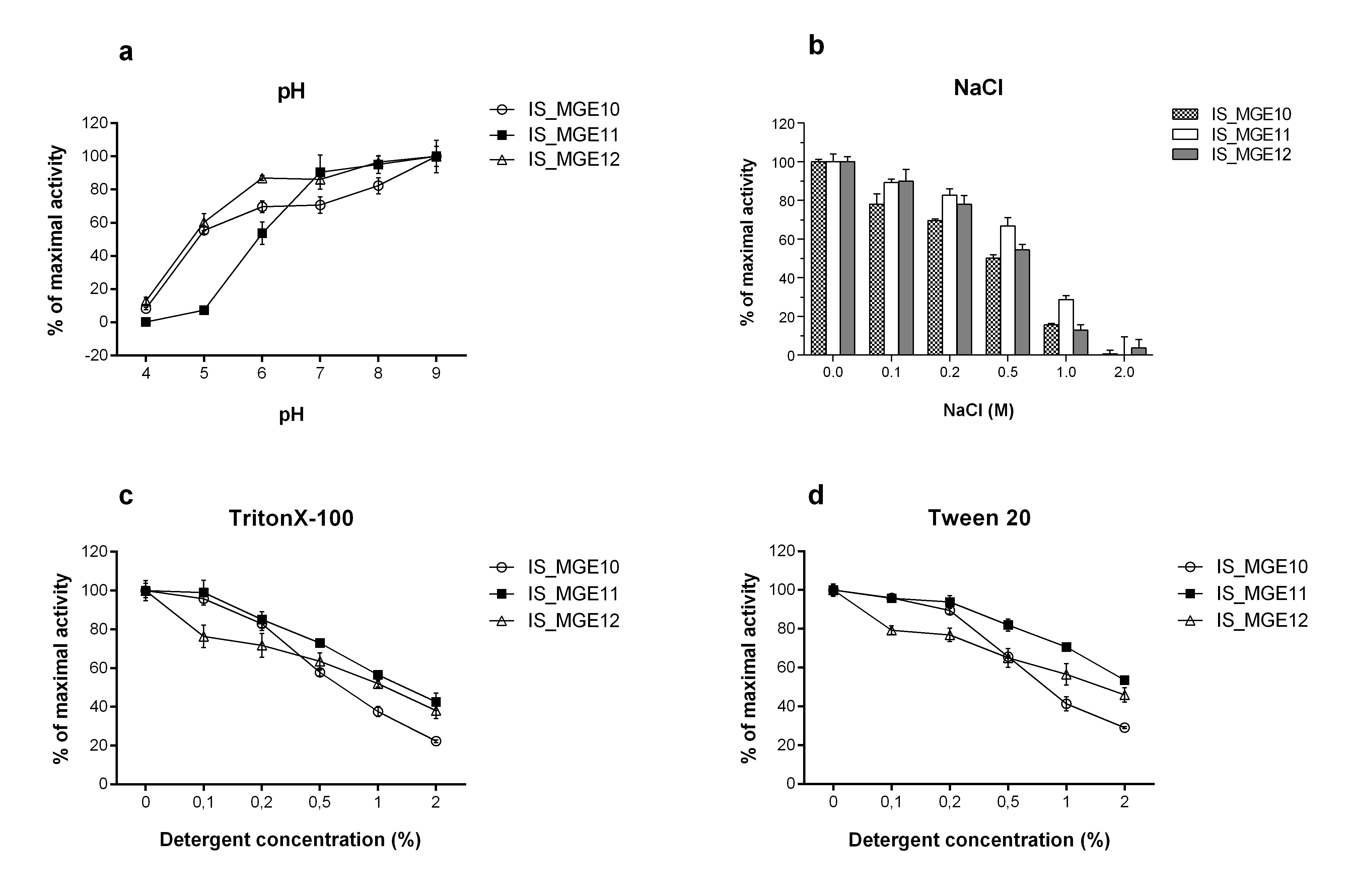

**Figure S4.** Carboxylesterase activity of IS10, IS11, and IS12: effect of pH, NaCl, and detergents. Esterase activity of purified proteins was measured as indicated using *p*NP-butyrate as substrate at 30ºC as described in Materials and Methods. Results are means ± SD from three independent experiments.

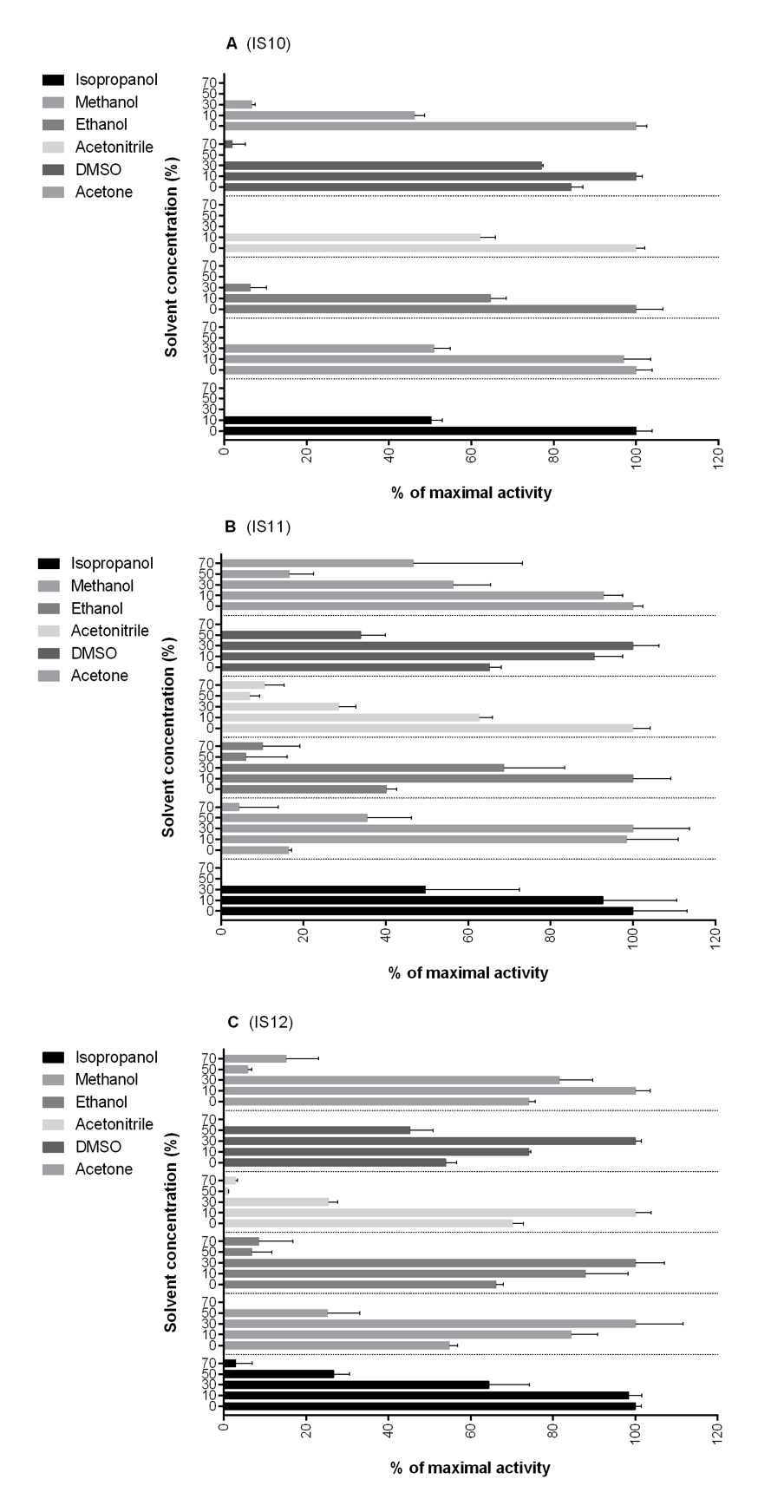

**Figure S5.** Effect of organic solvents on the activity of IS10, IS11 and IS12. Carboxylesterase activity of purified proteins was measured in the presence of increasing solvent concentrations (10, 30, 50, 70%, v/v) using *p*NP-butyrate as substrate at 30ºC as described in Materials and Methods. Results are means ± SD from three independent experiments.

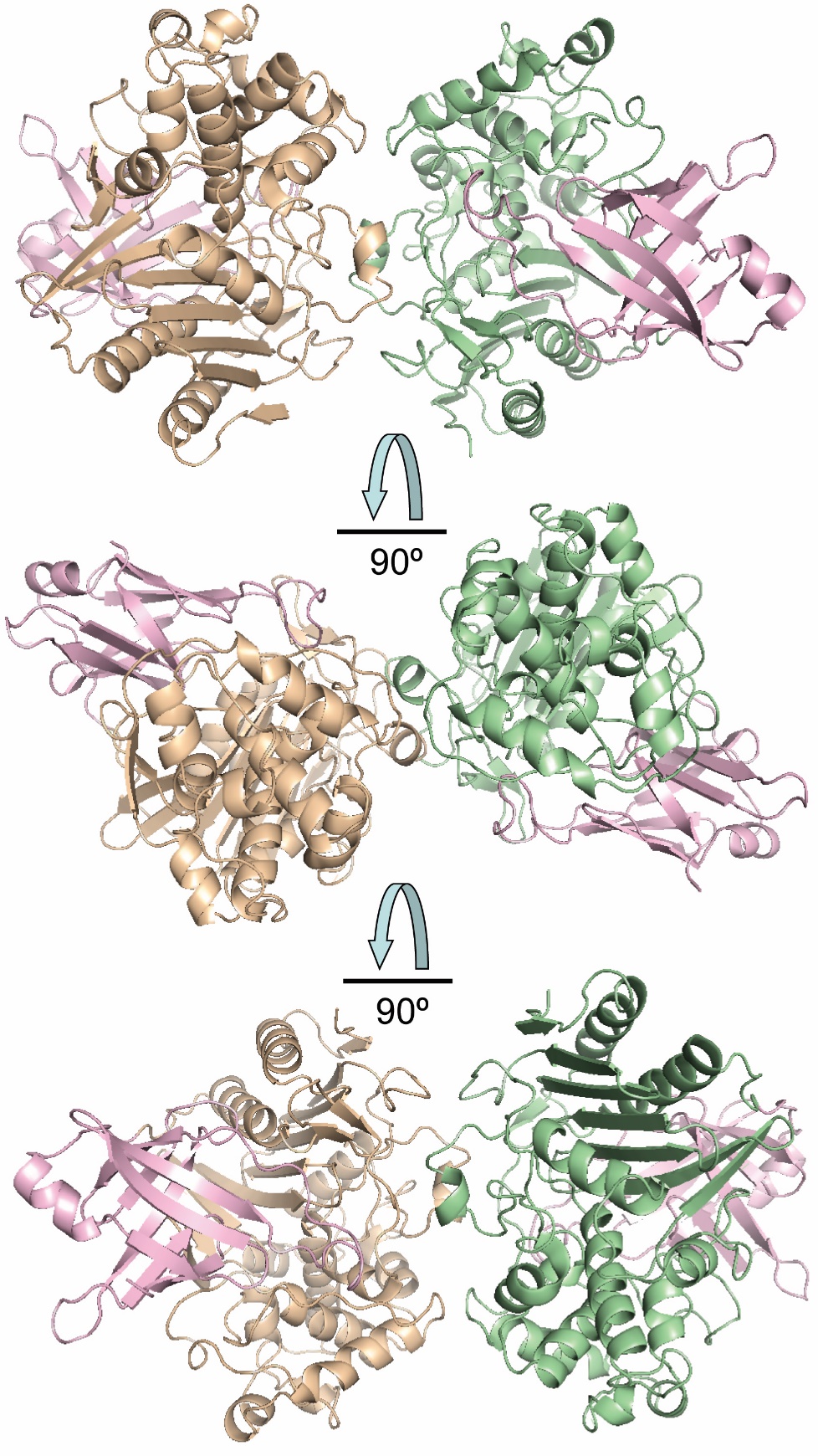
m

**Figure S6.** Overall crystal structure of IS11: three views of the IS11 dimer related by 90º rotations. The protein subunits are shown as ribbon diagrams with the core domains coloured wheat or green, whereas their lipocalin domains are coloured pale pink.

**Figure S7.** Oligomeric state of purified Ischia carboxylesterases. Elution profiles for purified proteins from size-exclusion chromatography using a Superdex 200 10/300 GL column. The numbers on the protein peaks indicate elution volumes (in ml), and corresponding molecular mass is reported (predicted monomer mass: 34.3 kDa for IS10, 49.4 kDa for IS11, and 33.9 kDa for IS12).

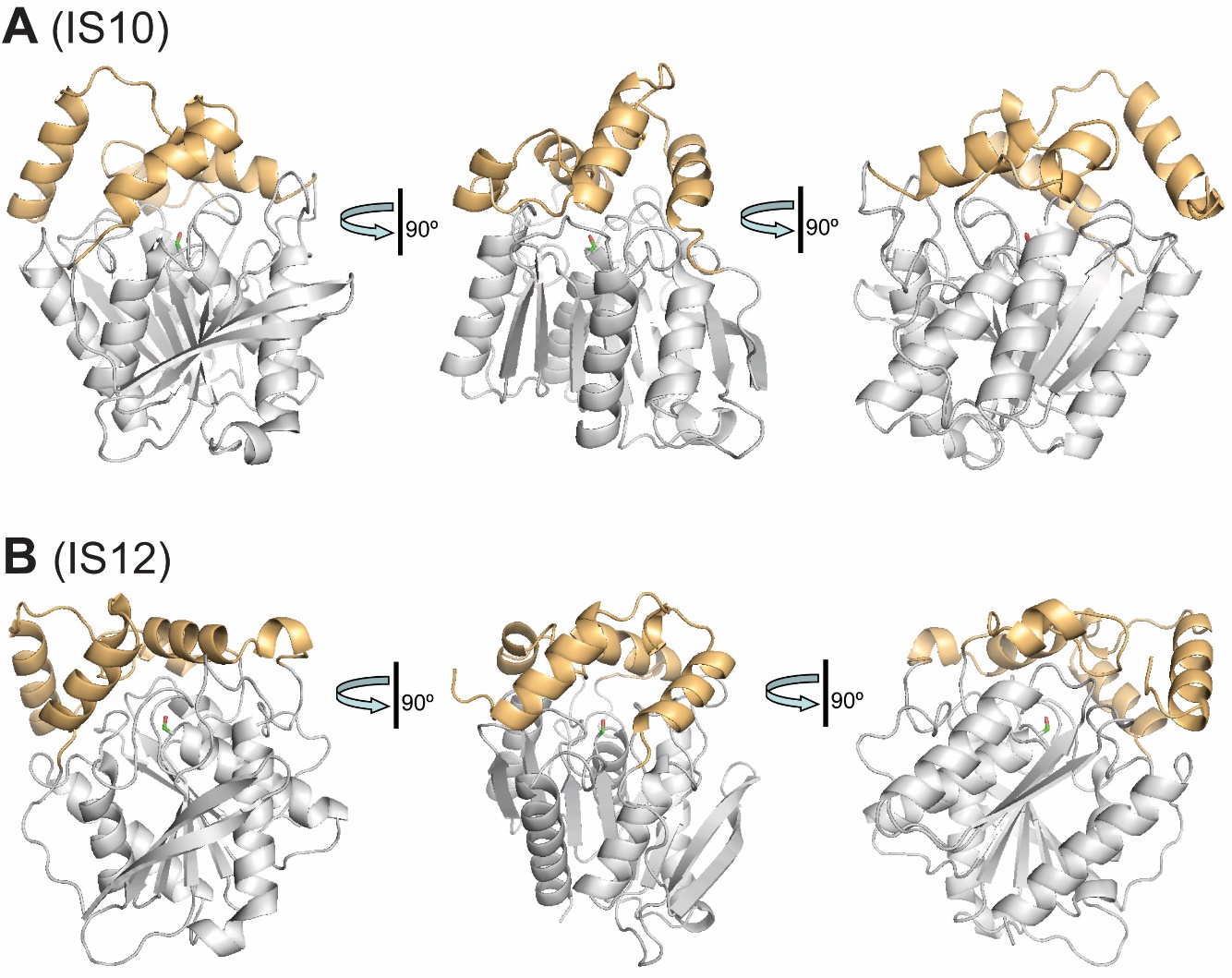

**Figure S8.** Structural models of IS10 (A) and IS12 (B) shown in three views related by a 90º rotation. Structural models were generated using the Phyre2 protein fold recognition server (1). Both models were constructed using the crystal structure of metagenomic carboxyl esterase Est8 (PDB code 4YPV (2) as the template (51% and 55% of sequence identity to IS10 and IS12, respectively; 100% confidence). The protein lid domain is coloured in light orange, whereas the core domain is shown in grey with the position of the active site indicated by the side chain of catalytic Ser155.

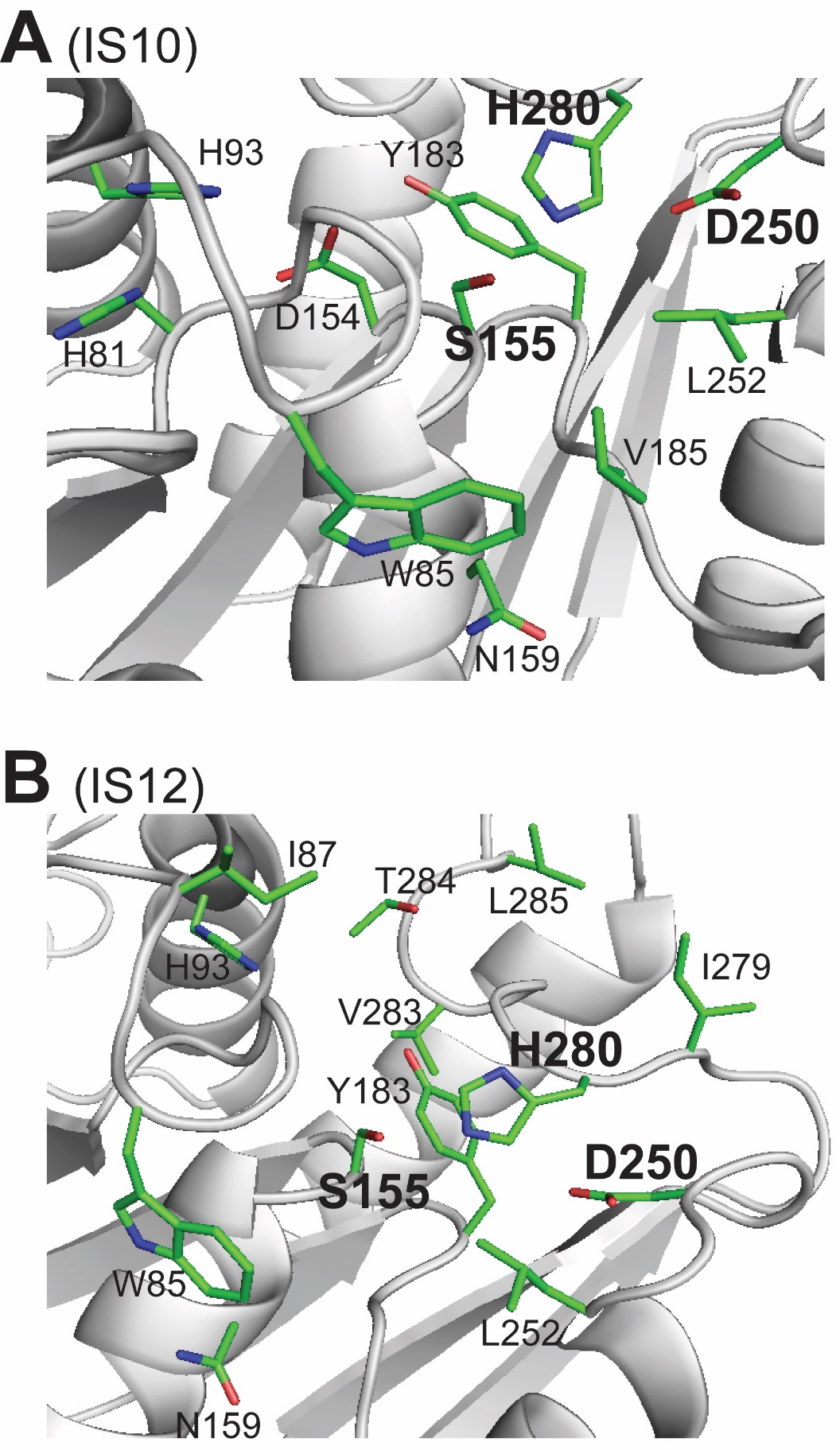

**Figure S9.** Structural models of IS10 and IS12: close-up view of the active sites. The protein ribbon is coloured in grey with amino acid side chains shown as sticks and carbon atoms coloured in green. The catalytic triad residues are indicated using larger labels (Ser155, His280, and Asp250), whereas other residues are potentially involved in substrate binding.

**Table S1.** Environmental (*in situ*) and experimental conditions for metagenome samples and enrichment cultures.

| **Sampling Site** | | | | **Enrichment Culture** | | | |
| --- | --- | --- | --- | --- | --- | --- | --- |
| **Name** | **ID** | **Temp** | **pH** | **Medium** | **pH** | **Substrate** | **Temp** |
| Cavascura | IS1 | 45°C | 8.5 | DSMZ 1374 | 7.5 | 1. PHB 2. PLA 3. Blend | 50°C |
|  | IS2 | 55°C | 7.0 | DSMZ 1374 | 7.5 | 1. PHB 2. PLA 3. Blend | 50°C |
| Maronti Beach | IS3 | 75°C | 4.5 | DSMZ 88 | 4.5 | 1. PHB 2. PLA 3. Blend | 75°C |
|  | IS4 | 85°C | 5.0 | DSMZ 88 | 4.5 | 1. PHB 2. PLA 3. Blend | 75°C |

a Polyester substrates used in enrichment cultures: PHB, polyhydroxybutyrate; PLA, polylactic acid; Blend, commercial polyester blend.

**Table S2.** Esterase activities of purified Ischia carboxylesterases against various monoesters analysed using a pH-shift assay (see Materials and Methods for details).

| **Substrates** | **Activity**, U/mg (± SD) | | |
| --- | --- | --- | --- |
|  | **IS10** | **IS11** | **IS12** |
| 1. (1*R*)-(-)-Menthyl acetate | 23.2 ± 0.7 | n.d. | 29.6 ± 0.9 |
| 2. (1*S*)-(+)-Menthyl acetate | 28.3 ± 0.8 | n.d. | 31.3 ± 0.9 |
| 3. 1-Naphthyl acetate | 2,575.7 ± 64.4 | 14.2 ± 1.9 | 1,413.9 ± 0.4 |
| 4. 1-Naphthyl butyrate | 352.5 ± 10.3 | 28.3 ± 1.4 | 505.6 ± 0.9 |
| 5. (1*R*)-(+)-Neomenthyl acetate | 20.6 ± 0.6 | n.d. | 27.0 ± 0.8 |
| 6. (1*S*)-(+)-Neomenthyl acetate | 11.6 ± 0.3 | n.d. | 14.2 ± 0.4 |
| 7. 2,4-Dichlorobenzyl 2,4-dichlorobenzoate | 150.5 ± 4.5 | n.d. | 38.6 ± 1.2 |
| 8. 3-Methyl-3-buten-1-yl acetate | 10.3 ± 0.3 | 2.6 ± 0.1 | 102.9 ± 3.0 |
| 9. Benzyl (*R*)-(+)-2-hydroxy-3-phenylpropionate | 28.3 ± 0.8 | 393.7 ± 8.9 | 29.6 ± 11.9 |
| 10. Butyl acetate | 136.4 ± 0.5 | n.d. | 144.5 ± 0.5 |
| 11. Cyclohexyl butyrate | 402.7 ± 12.4 | n.d. | 379.5 ± 11.4 |
| 12. Ethyl acetoacetate | 15.4 ± 0.5 | 1.3 ± 0.1 | 16.7 ± 0.5 |
| 13. Ethyl 2-chlorobenzoate | 131.2 ± 4.0 | n.d. | 149.2 ± 4.6 |
| 14. Ethyl (*R*)-(+)-4-chloro-3-hydroxybutyrate | 3.9 ± 0.2 | 24.4 ± 0.3 | 7.7 ± 0.8 |
| 15. Ethyl (*S*)-(−)-4-chloro-3-hydroxybutyrate | 19.3 ± 0.7 | 14.2 ± 0.7 | 23.8 ± 0.5 |
| 16. Ethyl 2-ethylacetoacetate | 16.7 ± 0.5 | 20.6 ± 0.6 | 20.6 ± 0.6 |
| 17. (+)-Ethyl D-lactate | 18.0 ± 0.6 | 28.9 ± 0.9 | 28.3 ± 0.9 |
| 18. (−)-Ethyl L-lactate | 25.7 ± 0.7 | 6.4 ± 0.8 | 25.7 ± 0.2 |
| 19. Ethyl 2-methylacetoacetate | 27.0 ± 0.8 | 10.3 ± 0.3 | 29.7 ± 0.8 |
| 20. Ethyl 3-oxohexanoate | 25.7 ± 0.8 | 9.0 ± 0.3 | 27.0 ± 0.8 |
| 21. Ethyl propionylacetate | 26.1 ± 0.8 | 18.0 ± 0.5 | 28.3 ± 0.8 |
| 22. Geranyl acetate | 467.0 ± 12.6 | 27.0 ± 0.9 | 519.8 ± 14.4 |
| 23. Glucose pentaacetate | 284.3 ± 8.8 | n.d. | 328.1 ± 10.4 |
| 24. Glyceryl triacetate | 979.1 ± 27.6 | 9.0 ± 0.3 | 1,071.7 ± 3.0 |
| 25. Glyceryl tributyrate (tributyrin) | 1,100.0 ± 32.0 | 30.9 ± 1.0 | 1,558.0 ± 47.9 |
| 26. Glyceryl tripropionate | 3,429.9 ± 88.9 | 32.2 ± 5.4 | 2,950.1 ± 1.0 |
| 27. Hexyl acetate | 34.7 ± 1.0 | n.d.a | 1.3 ± 0.1 |
| 28. Methyl glycolate | 3.9 ± 0.1 | n.d. | 6.4 ± 0.2 |
| 29. (−)-Methyl (*R*)-3-hydroxyvalerate | 12.9 ± 0.4 | 6.4 ± 0.5 | 15.4 ± 0.3 |
| 30. (+)-Methyl (*S*)-3-hydroxyvalerate | 6.4 ± 0.2 | 20.6 ± 0.3 | 7.7 ± 0.6 |
| 31. Octyl acetate | 154.4 ± 4.4 | n.d. | 128.7 ± 3.8 |
| 32. n-Pentyl benzoate | 16.7 ± 0.5 | n.d. | 14.4 ± 0.4 |
| 33. Phenyl acetate | 10,455.8 ± 15.7 | 21.9 ± 0.6 | 3,374.6 ± 15.4 |
| 34. Phenyl propionate | 8,792.3 ± 41.7 | 25.7 ± 0.8 | 8,219.8 ± 43.0 |
| 35. Phthalic acid diethyl ester | 29.6 ± 0.9 | n.d. | 105.5 ± 3.2 |
| 36. Propyl acetate | 20.6 ± 0.7 | n.d. | 25.7 ± 0.6 |
| 37. Propyl butyrate | 34.7 ± 0.1 | n.d. | 50.2 ± 1.5 |
| 38. Propyl hexanoate | 245.7 ± 7.0 | n.d. | 304.9 ± 8.8 |
| 39. $ϒ$-Valerolactone | 172.4 ± 5.4 | 60.5 ± 2.4 | 218.7 ± 6.7 |
| 40. Vinyl acetate | 74.6 ± 0.2 | n.d. | 66.9 ± 0.1 |
| 41. Vinyl benzoate | 1,100.0 ± 32.8 | n.d. | 1,084.6 ± 32.0 |
| 42. Vinyl butyrate | 200.7 ± 5.7 | n.d. | 616.3 ± 18.3 |
| 43. Vinyl crotonate | 5.1 ± 0.2 | n.d. | 90.1 ± 2.7 |
| 44. Vinyl propionate | 187.8 ± 5.5 | n.d. | 833.7 ± 24.0 |

a n.d., not detectable.

**Table S3.** X-ray crystallographic statistics for the structure of IS11.

| **PDB code** | **7SPN** |
| --- | --- |
| **Data collection** |  |
| Space group | P2 |
| Unit cell  *a*, *b, c* (Å)  α, β, γ, (°) | 50.22, 103.36, 94.86  90, 126.56, 90 |
| Resolution, Å | 50.00 – 2.92 |
| R*_merge_*a  R*_pim_*b | 0.115 (0.831)*  0.069 (0.528) |
| CC_1/2_ | 0.993 (0.572) |
| *I* / σ(*I)* | 11.96 (1.10) |
| Completeness, % | 98.4 (97.6) |
| Redundancy | 3.6 (3.2) |
| **Refinement** |  |
| Resolution, Å | 45.09 – 2.92 |
| No. unique reflections:  working, test | 18739, 1834 |
| *R*-factor/free *R­*-factorc | 20.7/26.0 (30.8/38.2) |
| No. refined atoms, molecules  Protein  Water | 6990, 2  85 |
| *B*-factors  Protein  Water | 71.3  64.0 |
| r.m.s.d.  Bond lengths, Å  Bond angles, ° | 0.002  0.534 |

*values in brackets refer to highest resolution shells.

a*R*_merge_ = Σ_hkl_Σ_j_|*I*_hkl.j_ - 〈*I*_hkl_〉|/Σ_hkl_Σ_j_I_hk,j_, where *I*_hkl,j_ and 〈*I*_hkl_〉 are the *j*th and mean measurement of the intensity of reflection *j*.

b*R*_pim_ = Σ_hkl_√(n/n-1) Σn_j=1_|*I*_hkl.j_ - 〈*I*_hkl_〉|/Σ_hkl_Σ_j_I_hk,j_

c*R* = Σ|F_p_obs – F_p_calc|/ΣF_p_obs, where F_p_obs and F_p_calc are the observed and calculated structure factor amplitudes, respectively.

**Supplementary References**

1. Kelley, L. A., Mezulis, S., Yates, C. M., Wass, M. N., Sternberg M. J. (2015) The Phyre2 web portal for protein modeling, prediction and analysis. *Nat Protoc* **10,** 845-858.
2. Pereira, M. R., Maester, T. C, Mercaldi, G. F., de Macedo Lemos, E. G., Hyvönen, M., Balan, A. (2017) From a metagenomic source to a high-resolution structure of a novel alkaline esterase. *Appl Microbiol Biotechnol* **101**, 4935-4949.
